# Benchmarking gene expression reconstruction from single-cell latent representations

**DOI:** 10.64898/2026.06.15.731445

**Authors:** Xiaotong Fu, Dominik Klein, Egor Antipov, Alessandro Palma, Alejandro Tejada-Lapuerta, Mojtaba Bahrami, Louis B. Kümmerle, Manuel Lubetzki, Francesco Paolo Casale, Malte D. Luecken, Fabian J. Theis

## Abstract

Single-cell transcriptomics is typically modeled in low-dimensional latent representations that improve the signal-to-noise ratio of the data. Such representations underpin data integration, cell state discovery, and perturbation prediction, with applications ranging from large-scale organ atlases to latent trajectory modeling. Recent virtual cell approaches further leverage these representations to predict cellular responses as distributional shifts in latent space. Each of these applications ultimately requires faithful gene expression reconstruction from latent spaces for biological interpretation, enabling gene-level analysis of predicted perturbed or batch-corrected cells. Yet representation choice is typically treated as an implementation detail rather than a primary modeling decision, with no systematic evaluation of how well latent representations support gene expression reconstruction. Here, we introduce ReconEval, a benchmark for evaluating gene expression reconstruction from single-cell latent spaces. We benchmark two classes of latent representations: end-to-end trained models such as PCA, autoencoders, and variational autoencoders, and pretrained single-cell foundation model embeddings coupled to newly trained decoders. Reconstruction is evaluated both directly and after latent-space perturbation prediction. Across perturbational and observational datasets totaling over 100 million cells, our metric suite quantifies statistical fidelity; biological signal preservation, including differential expression, coexpression, cell-cycle structure, cytokine response and pathway activity; and perturbation-specific effects. We find that autoencoders achieve the strongest stand-alone reconstruction at low dimensionality, while variational regularization does not improve generalization in reconstruction. Frozen foundation model embeddings retain recoverable gene-level information, with reconstruction quality depending strongly on decoder architecture and pretraining objective. In latent perturbation modeling, high-dimensional PCA matches foundation model embeddings, while low-dimensional AE embeddings are optimal for flow-based generative models. Overall, reconstruction depends critically on the interplay between representation and downstream model, and simpler representations can outperform complex alternatives given appropriate capacity. Our benchmark establishes reconstruction as a critical evaluation axis for single-cell foundation models. We envision it improving the biological interpretability of latent-space modeling, a prerequisite for future virtual cell models to be validated by domain experts and grounded in biology.

## Introduction

Owing to its transcriptome-wide readout at an unprecedented resolution, the single-cell RNA-seq (scRNA-seq) technology has transformed our understanding of cellular biology, such as state heterogeneity, disease programs, and cellular dynamics^1^. However, scRNA-seq data are sparse, noisy, and high-dimensional, making many downstream analyses and modeling tasks challenging when performed directly in gene space^2^. Lower-dimensional representations have therefore become the computational backbone of single-cell analysis: they underpin cell-type annotation and rare-population discovery^3^, enable atlas-scale integration across studies, tissues, and technologies^4,5^, and often form the basis for perturbation modeling, where shifts in the latent space are decoded back to gene expression to predict cellular responses to interventions^6^.

Recently, a central emerging goal in single-cell modeling is the “virtual cell”: predictive models that simulate how a cell’s transcriptome changes under external interventions or internal processes, including drug treatment, cytokine stimulation, or disease progression^7^. Many state-of-the-art perturbation methods achieve this by modeling shifts in a latent cell space and decoding the shifted representations back to gene expression, a paradigm we refer to as latent shift modeling^8–14^. For example, CellFlow uses generative flow matching^15^ to predict perturbation responses in compressed representations^16^. Meanwhile, STATE utilizes a transformer backbone to minimize the energy distance between predicted and true cell populations^17^.

In practice, single-cell workflows converge on a small number of latent representations^2^. PCA remains the default linear embedding in standard analysis pipelines, while variational autoencoders (VAEs), most prominently scVI^4^ and extensions thereof^18,19^, have become the prevailing nonlinear framework, adopted by the majority of perturbation prediction methods^20^ and atlas integration studies^21,22^. Despite this widespread adoption, the representation for reconstruction is typically treated as a fixed implementation choice rather than a modeling decision that is explicitly validated^16,23^. This is consequential because in latent shift modeling, a downstream model, such as a perturbation prediction model, operates directly in the latent space. If a latent space does not retain sufficient information for faithful gene expression reconstruction, decoded cells may be biologically implausible even when the latent transformation is well fit. This is of particular importance because the biological characterization of any cell, including differential expression, pathway analysis, and gene-gene coexpression structure is ultimately performed in gene expression space rather than in abstract latent coordinates.

The recent emergence of single-cell foundation models adds a new dimension to this representation choice: pretrained encoders based on metric learning^24^, contrastive objectives^25^, or transformer architectures^17,26^ provide general-purpose embeddings trained on tens to hundreds of millions of cells, useful across discriminative and retrieval tasks. Yet many of these models are decoder-free and rarely evaluated for gene-level reconstruction. It is therefore unclear how much gene-resolution information these embeddings retain, and whether they can serve as embedding spaces for latent shift modeling when coupled with a learned decoder. Given the central role of representation learning in single-cell data analysis, a comprehensive comparison of embedding methods with respect to gene expression reconstruction is essential. A key challenge in evaluating reconstruction is that no established framework exists for systematically assessing whether decoded gene expression retains biologically meaningful information: existing benchmarks of latent representations have evaluated data integration^5,27,28^ or perturbation prediction^17,29^, but reconstruction quality remains unaddressed across the full range of latent representation families and downstream models.

Here, we present ReconEval, a benchmark of gene expression reconstruction across latent representation models, including PCA, AEs, and VAEs, as well as pretrained foundation model embeddings coupled with trained decoders (Fig. 1a). The latter setting remains largely unexplored despite its direct relevance to how foundation models are deployed. Reconstruction is essential for latent shift modeling, in which perturbation-induced shifts (e.g., from drug treatment, cytokine stimulation, viral infection, cell division, or batch effects^5,30–33^) are decoded back to gene-level changes (Fig. 1b). To this end, we consider three reconstruction tasks across two settings: stand-alone reconstruction and latent shift reconstruction (Fig. 1c). Stand-alone reconstruction comprises (i) end-to-end reconstruction, where encoder and decoder are jointly optimized, and (ii) foundation-model reconstruction, where a pretrained foundation model serves as a frozen encoder and a dedicated decoder is trained separately to map fixed embeddings back to gene expression. Latent shift reconstruction reflects the practical use case motivating this benchmark: (iii) a perturbation prediction model operates on the embedding before decoding, allowing us to assess whether reconstruction quality is maintained after latent-space manipulation. To evaluate these tasks, we develop a unified metric suite spanning statistical fidelity, biology-informed measures, and perturbation effect retention. We evaluate reconstruction across three datasets spanning perturbational and observational regimes at different scales: Tahoe-100M^34^, Human cytokine dictionary (PBMC-10M)^35^, and the Human Lung Cancer Atlas (LuCA)^36^. Our benchmark formalizes reconstruction as a testable property of both trained latent spaces and pretrained foundation model embeddings, provides practical guidance for representation selection, and reveals that optimal latent-space design depends critically on the interplay between the representation and the downstream model.

**Fig. 1.**
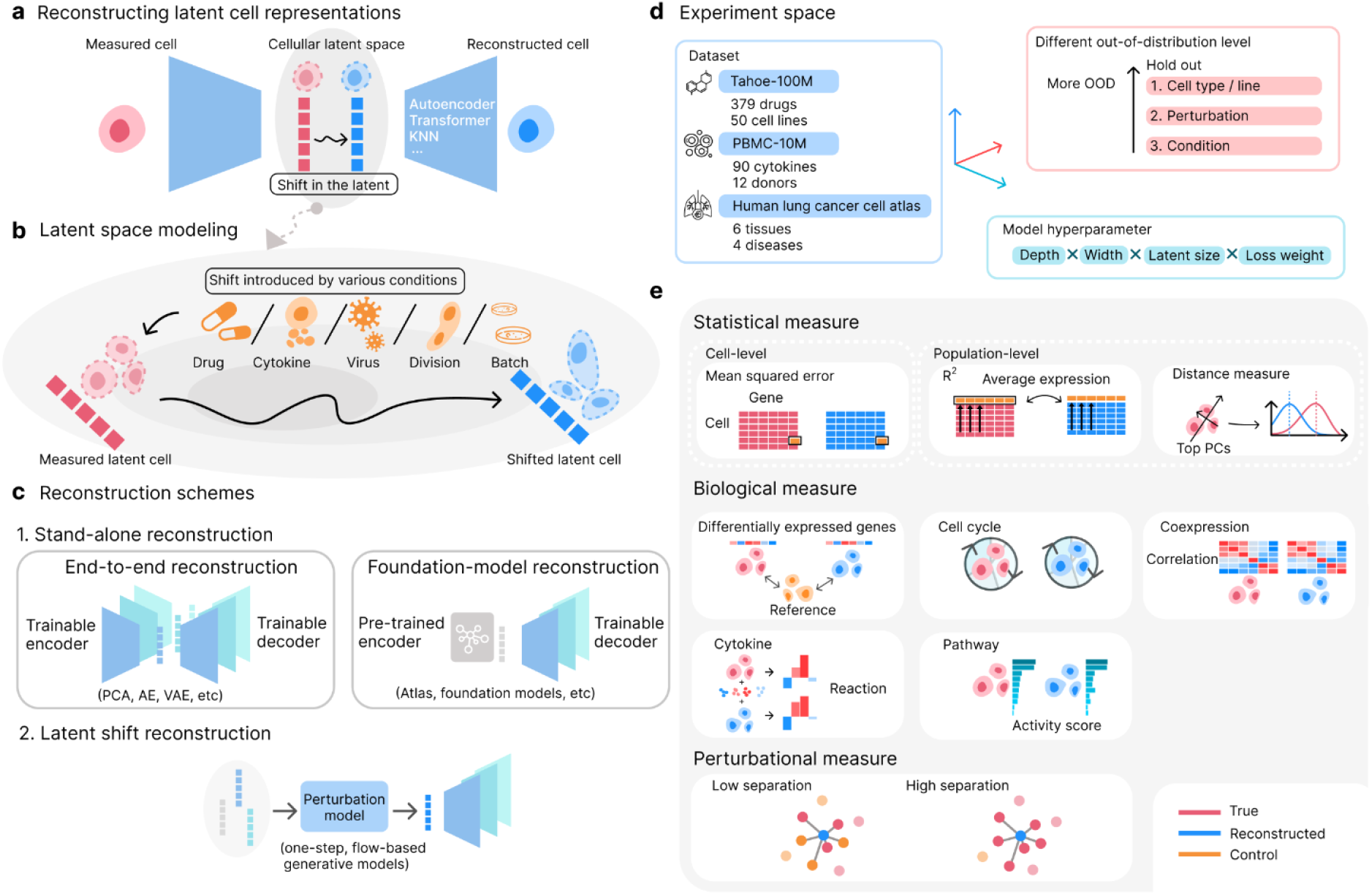
Overview of ReconEval: a benchmark for gene expression reconstruction from single-cell latent representations. **a**, Reconstruction maps a latent cell representation back to a gene expression profile via a decoder. **b**, Reconstruction is essential for latent shift modeling, in which perturbation-induced shifts are decoded back to gene-level changes. **c**, Three reconstruction paradigms are evaluated across two perspectives. (1) Stand-alone reconstruction: end-to-end reconstruction (left), in which encoder and decoder are trained jointly (PCA, AE, VAE), and foundation-model reconstruction (right), in which a pretrained, frozen encoder provides fixed embeddings to a separately trained decoder (SE embedding from STATE, scConcept, scGPT, SCimilarity). (2) Latent shift reconstruction: reconstruction quality is assessed after a latent shift prediction model (STATE, CellFlow) operates on the embeddings. **d**, The experiment space spans three datasets (Tahoe-100M, PBMC-10M, LuCA), three levels of out-of-distribution generalization (cell type/line, perturbation/donor, and condition-level hold-out), and a systematic hyperparameter grid covering model depth, width, latent dimensionality, and loss weight. **e**, Reconstruction quality is assessed using metrics spanning three categories: statistical fidelity (cell-level MSE, population-level R^2^, energy distance), biological signal preservation (differentially expressed gene recovery, cell-cycle composition, gene-gene coexpression, cytokine reaction, pathway activity), and perturbational effect retention (perturbation separation).

## Results

### Benchmarking gene expression reconstruction across single-cell representations

Gene expression reconstruction from latent representations is essential whenever biological interpretation is required, yet its quality has not been systematically evaluated. This problem becomes even more pressing given the rapid evolution of representation learning and latent shift modeling in single-cell genomics. We therefore benchmark reconstruction performance across representations and downstream models. We evaluate different representation families in three reconstruction tasks across two perspectives (Fig. 1c): In *end-to-end reconstruction*, we compare PCA, AEs, and VAEs, which are widely used in single-cell genomics and machine learning more broadly. In *foundation-model reconstruction*, we evaluate embeddings learned from four foundation models (SE embedding from STATE^1^, scConcept, scGPT, SCimilarity)^17,24–26^ paired with three decoder architectures (multilayer perceptron (MLP), k-nearest neighbors (KNN), and Transformer). This separation mirrors how embeddings are actually obtained in practice: classical representation models are trained on the dataset of interest, whereas foundation model embeddings are typically used as frozen, general-purpose representations with a small task-specific head. In *latent shift reconstruction*, we assess whether reconstruction quality is maintained after a perturbation prediction model operates on the embedding. To capture the diversity of current latent shift approaches, we evaluate two distinct paradigms for learning distributional shifts in latent space: flow-based generative models, which learn continuous paths from unperturbed to perturbed cell distributions^8,10,11,16^, and one-step generative models which minimize the distance between predicted and true perturbed distributions^17,23^. We evaluate STATE as an instance of one-step generative models and CellFlow as an instance of flow-based generative models, providing complementary perspectives on how representation properties affect downstream tasks.

To probe generalization across biological regimes, we benchmark across three scRNA-seq datasets (Fig. 1d): Tahoe-100M (drug perturbations, ∼89M cells)^34^, Parse PBMC-10M^35^ (cytokine perturbations, ∼10M cells), and the Single-Cell Lung Cancer Atlas^36^ (LuCA; observational, ∼900K cells). Together, these datasets cover distinct perturbation modalities (drug, cytokine, none), biological systems (cell lines, blood cells, lung cancer), and scales, while providing substantial condition diversity (See Methods, Table 1). For each dataset, we define multiple train–test splits at increasing out-of-distribution (OOD) difficulty and perform pre-registered hyperparameter searches for each model family (see Methods, Supplementary Table 1, Supplementary Table 2). OOD difficulty is quantified in Extended Data Fig. 1, where PBMC-10M and LuCA show substantially higher heterogeneity than Tahoe-100M. Since latent dimensionality is a primary bottleneck for reconstruction and directly determines the capacity available for downstream latent-space modeling, we benchmark models across latent dimensionalities d ∈ {10,32,128,512,2048}. Preprocessing and split construction details are provided in Methods.

**Table 1.**
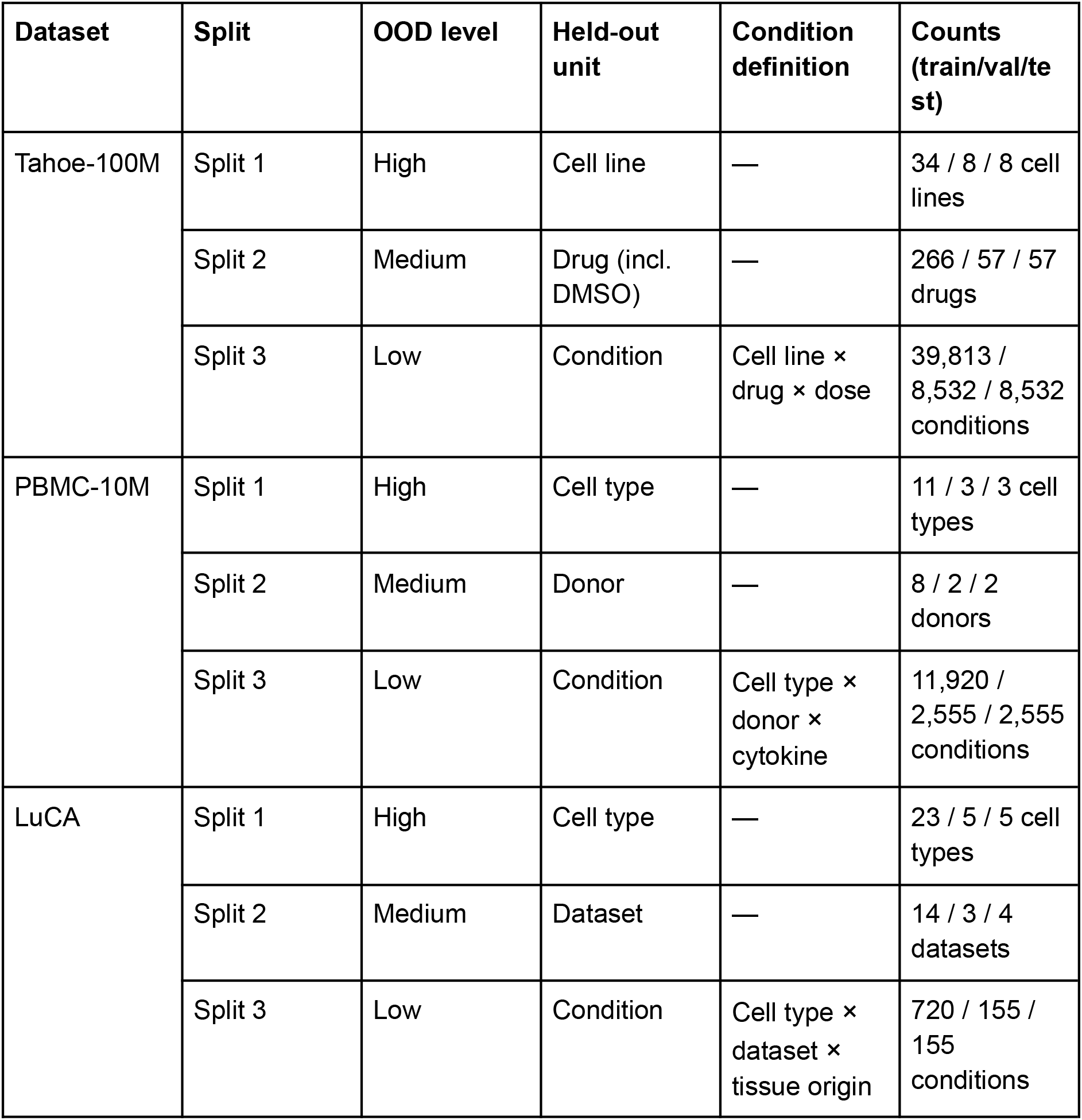
Train/validation/test splits for each dataset at three OOD difficulty levels. For each dataset, we constructed three splits with decreasing OOD difficulty by holding out different covariate units. Split 1 is the most OOD (holding out cell types or cell lines), Split 2 is intermediate (holding out datasets, donors, or drugs), and Split 3 is the least OOD (holding out specific condition combinations). Counts are reported as train/validation/test at the level of the held-out unit. Conditions correspond to unique combinations of the listed covariates.

| Dataset | Split | OOD level | Held-out unit | Condition definition | Counts (train/val/test) |
| --- | --- | --- | --- | --- | --- |
| Tahoe-100M | Split 1 | High | Cell line | — | 34 / 8 / 8 cell lines |
|  | Split 2 | Medium | Drug (incl. DMSO) | — | 266 / 57 / 57 drugs |
| | Split 3 | Low | Condition | Cell line × drug × dose | 39,813 / 8,532 / 8,532 conditions |
| PBMC-10M | Split 1 | High | Cell type | — | 11 / 3 / 3 cell types |
|  | Split 2 | Medium | Donor | — | 8 / 2 / 2 donors |
| | Split 3 | Low | Condition | Cell type × donor × cytokine | 11,920 / 2,555 / 2,555 conditions |
| LuCA | Split 1 | High | Cell type | — | 23 / 5 / 5 cell types |
|  | Split 2 | Medium | Dataset | — | 14 / 3 / 4 datasets |
| | Split 3 | Low | Condition | Cell type × dataset × tissue origin | 720 / 155 / 155 conditions |

We evaluate reconstruction with metrics across three axes (Fig. 1e). Statistical measures capture numerical fidelity at both the single-cell level (mean squared error, *MSE*) and the population level (*R*^*2*^; energy distance, *Energy dist*.), accounting for the stochasticity inherent in single-cell measurements. Biological measures quantify whether reconstructions preserve signals relevant to common downstream analyses: recovery of differentially expressed genes (*DEG dice@100*) and gene-level effect sizes (log-fold-change correlation, *LogFC Sp*.), consistency of biological programs through pathway activity (*Pathway*) and cytokine response (*Cytokine*), and preservation of broader biological structure via gene-gene coexpression (*Coexpression*) and cell-cycle composition (*Cell cycle)*. A perturbational measure (*KNN purity*) assesses perturbation-specific effects in gene expression space by quantifying the separability of reconstructed perturbed cells from control cells relative to true perturbed cells. All metrics are computed per condition, defined as a unique combination of cell identity and treatment context (e.g., cell type × drug × dose), and reported on a held-out test set of conditions. Together, these metrics provide a unified evaluation suite bridging statistical fidelity, biological interpretability, and perturbation effect retention. We additionally compute an overall rank-percentile score, enabling consistent comparison across heterogeneous measures. See Methods for detailed definitions and formulas.

### AEs achieve the strongest stand-alone reconstruction at low latent dimensionality

We first evaluated end-to-end reconstruction, the most widely used representation learning scheme in single-cell genomics, in which both encoder and decoder are trained jointly. We compared three architecture families (PCA, AE, and VAE) across all three datasets and OOD splits. For AE and VAE models, we additionally assessed three library-size handling strategies: no explicit modeling, learned modeling, and observed library size as input (see Methods). Unless otherwise noted, we present results at d = 128, which avoids both severely bottlenecked and near-saturated regimes and provides a stable comparison point across models (See Extended Data Fig. 2). A full performance overview is provided in Extended Data Fig. 3.

Aggregating rank-percentile scores across all dataset–split combinations at d = 128 (Fig. 2a), AEs consistently achieved the highest overall reconstruction performance, outperforming both VAE and PCA on both statistical and biological measures. Meanwhile, varying library-size handling produced relatively modest changes compared to differences between architecture families: for AEs, modeled or observed library size slightly improved performance, whereas VAEs showed a performance drop under the modeled setting (Fig. 2a). The asymmetry likely reflects two competing influences: providing sequencing depth as additional biological signal can aid reconstruction, while jointly inferring it as a latent variable adds optimization complexity to the VAE objective. Based on this observation, we then opt for AE with observed library size and VAE with no library size as the best-performing configurations for subsequent analyses (Extended Data Fig. 3 shows the full comparison). The relative ordering of model families was robust under increasing OOD difficulty and stable across datasets (Fig. 2b, top), though absolute reconstruction accuracy degraded for all models under more challenging OOD regimes (Extended Data Fig. 4). Notably, AEs maintained the strongest reconstruction even under the most challenging OOD regime. We further examined scaling behavior across latent dimensionalities (Fig. 2b, bottom, Extended Data Fig. 5): AEs and VAEs compressed information effectively into representations as low as 10 dimensions with only modest gains at higher dimensionalities, suggesting that nonlinear models extract most reconstructive signal even under severe compression. AEs outperformed VAEs at all dimensions, while PCA showed steeper scaling, starting substantially lower but matching or exceeding nonlinear models at d = 2,048.

**Fig. 2.**
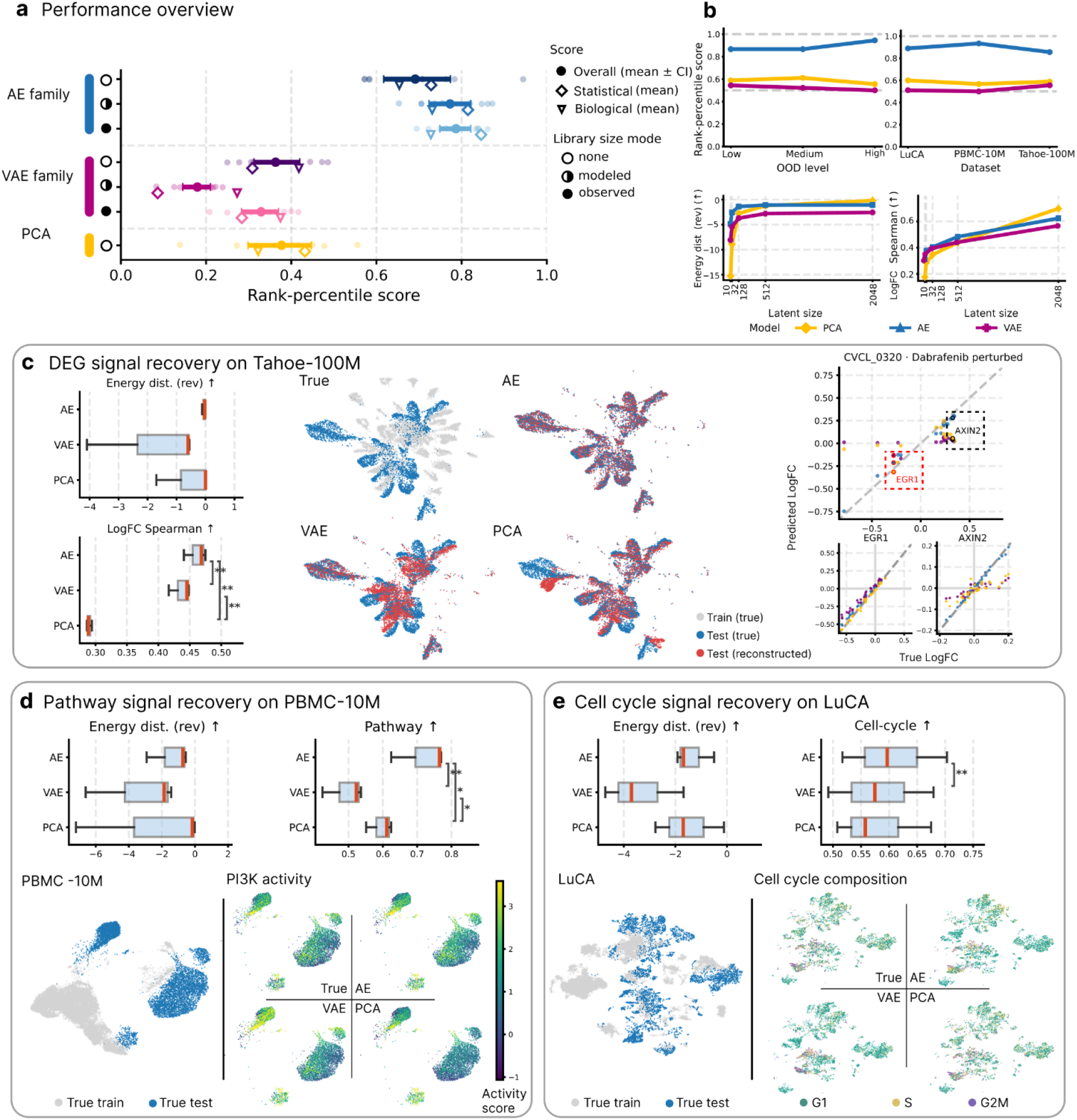
Autoencoders outperform alternatives across model families, datasets, and out-of-distribution regimes in end-to-end reconstruction. **a**, Performance overview. Each model configuration is characterized by its architecture (AE, VAE, PCA) and library-size mode (none, modeled, observed). For every dataset–split combination we compute rank-percentile scores for statistical fidelity (◇) and biological signal preservation (▽); the overall score (◆, mean ± 95% Confidence Interval) is their equally weighted average. **b**, Robustness to distribution shift and dataset identity. Top left: overall rank-percentile score as a function of OOD level. Top right: overall rank-percentile score across datasets. Bottom: metric scaling with latent dimensionality on LuCA, illustrated by Energy distance (statistical, reversed to be larger better) and LogFC Spearman correlation (biological). Within each architecture family, the best-performing library-size configuration is shown. **c**, Reconstruction quality on Tahoe-100M. Left: Energy distance (reversed) and LogFC Spearman across experiments, with significance markers from paired t-tests of conditions (*p < 0.05, **p < 0.01, ***p < 0.001). Middle: UMAP overlays of true training cells, true test cells, and reconstructed test cells. Right: predicted versus true log-fold-change for an example perturbation (Dabrafenib on cell line CVCL_0320), with insets highlighting marker genes *EGR1* and *AXIN2*. **d**, Pathway signal recovery on PBMC-10M. Top: Energy distance (reversed) and Pathway score across experiments, with significance markers as in **c**. Bottom: UMAPs showing true training and test cells (left) and pathway activity overlaid on test cells for ground truth, AE, VAE, and PCA reconstructions (right, *PI3K* shown as an example). **e**, Cell-cycle signal recovery on LuCA. Top: Energy distance (reversed) and Cell-cycle score across experiments, with significance markers as in **c**. Bottom: UMAPs showing true training and test cells (left) and cell-cycle phase composition (G1, S, G2/M) for ground truth, AE, VAE, and PCA reconstructions (right).

To examine reconstruction quality through the lens of dataset-specific biology, we selected a biologically motivated metric for each dataset and focused on the most challenging OOD split (holding out entire cell types or lines). On Tahoe-100M (Fig. 2c), a large-scale drug perturbation atlas, we focused on recovery of perturbation-induced transcriptional changes. AE achieved the highest *R*^*2*^ and lowest *MSE*, and most faithfully recovered log-fold-change structure among differentially expressed genes across drugs. Predicted versus true log-fold changes for a representative perturbation (Dabrafenib on CVCL_0320) illustrate that AE captured both global effect sizes and individual marker-gene shifts (e.g., *EGR1, AXIN2*), whereas VAE and PCA showed attenuated or noisier recovery (Fig. 2c, right). On PBMC-10M (Fig. 2d), a cytokine perturbation dataset in which treatment effects manifest as coordinated transcriptional programs, we focused on pathway activity recovery. AEs yielded significantly higher pathway activity correlations than VAE and PCA, with PCA outperforming VAE. We illustrate this with *PI3K*, a pathway central to cytokine receptor signaling and immune cell activation; qualitative comparison of *PI3K* activity scores on held-out cell types confirmed that AE reconstructions most closely recapitulated the distribution of pathway activation observed in ground-truth data. On LuCA (Fig. 2e), an observational patient tissue atlas, we focused on preservation of cell-cycle composition. AEs achieved higher cell-cycle phase retention than both VAE and PCA, which showed comparable performance to each other.

Across all three datasets, out-of-distribution levels, and latent dimensionalities, AEs provided the strongest end-to-end reconstruction, with consistent gains on both statistical and biological measures, including on held-out cell types and cell lines not seen during training. The additional regularization imposed by the variational prior in VAEs did not translate into improved reconstruction even under the most challenging out-of-distribution conditions. PCA, despite its linearity, proved competitive at higher latent dimensionalities, approaching AE-level fidelity when given sufficient capacity.

### Pretrained foundation model embeddings retain recoverable gene expression signal

We next evaluated foundation-model reconstruction, a setting that has remained mostly unexplored: a pretrained foundation model encoder is kept frozen and only a decoder is trained to map its embeddings back to gene expression (Methods). We evaluated embeddings learned from four foundation models representing distinct modeling strategies (SE embedding from STATE, scConcept, scGPT, and SCimilarity). To assess how reconstruction quality depends on both the foundation model embeddings and decoder architecture, we paired each embedding with three decoder architectures (MLP, KNN, and Transformer; see Methods). To avoid data leakage and better reflect genuine transfer usage, we report results on PBMC-10M, which was not included in the pretraining data of any evaluated model. UMAP projections of the pretrained embeddings (Fig. 3a) reveal that all four models capture broad cell-type structure in PBMC-10M, but differ in the granularity and separation of subpopulations.

**Fig. 3.**
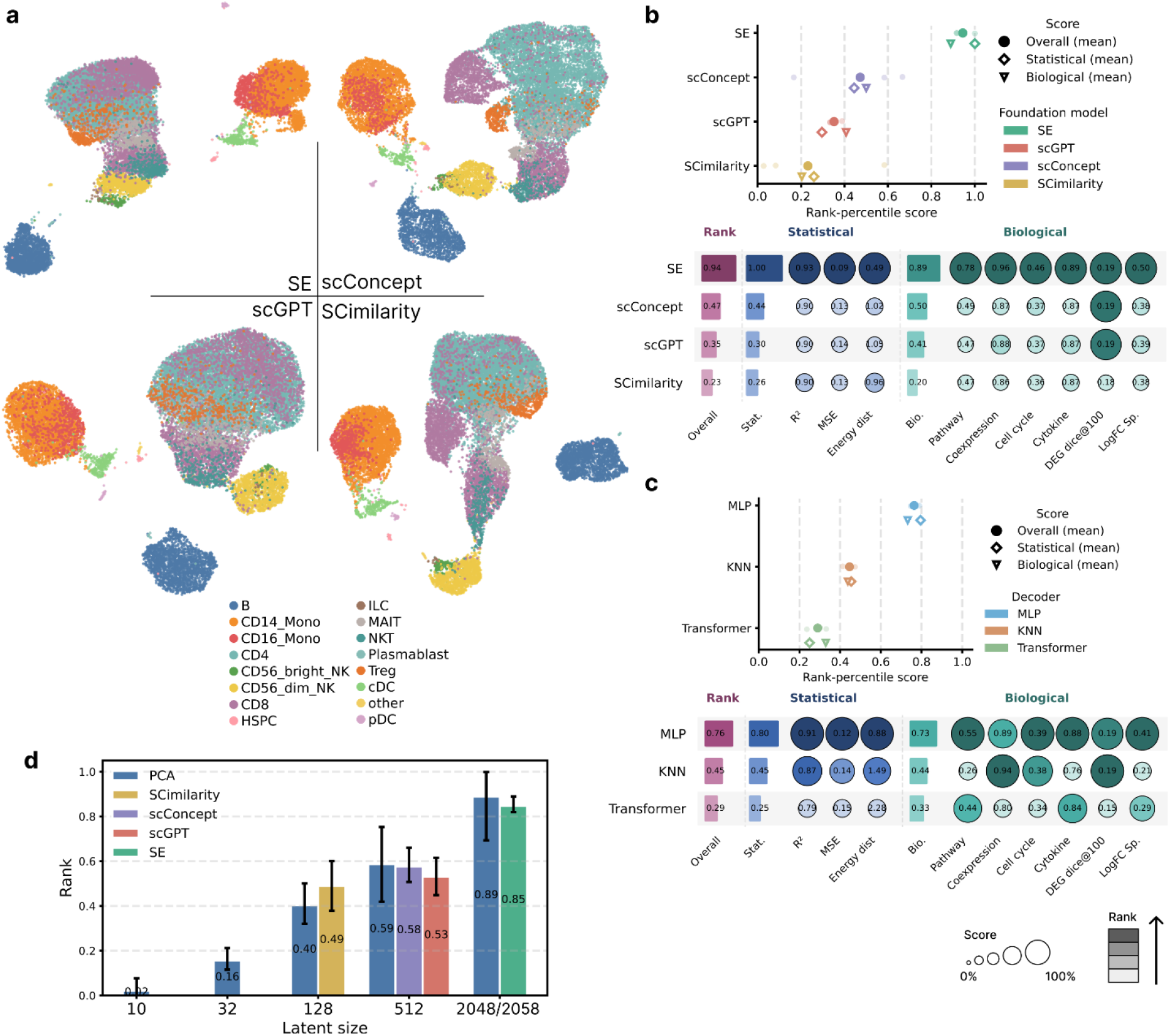
Foundation model embeddings retain recoverable gene-level information through trained decoders in foundation-model reconstruction. **a**, Latent-space structure of pretrained foundation model embeddings. UMAP projections of PBMC-10M cells colored by cell type for SE, scConcept, scGPT, and SCimilarity. **b**, Reconstruction performance of foundation model embeddings paired with MLP decoders. Top: for each foundation model, rank-percentile scores for distributional fidelity (◇) and biological signal preservation (▽); the overall score (◆) is their equally weighted average. Means of each split are shown in the background. Bottom: per-metric breakdown of statistical and biological measures. **c**, Decoder architecture effect on reconstruction quality. Top: performance overview for MLP, KNN, and Transformer decoders, aggregated across all four foundation models and three dataset splits. Bottom: per-metric breakdown as in **b. d**, Comparison of foundation model embeddings with PCA at matched dimensionality. Bars show overall rank-percentile score; error bars indicate variability across individual metrics.

Comparing foundation model embeddings with MLP decoders (Fig. 3b, top), SE achieved the highest overall reconstruction performance, followed by scConcept, scGPT, and SCimilarity. SE and scGPT showed relatively consistent performance across splits, whereas scConcept and SCimilarity exhibited higher variability. Performance generally degrades with increasing OOD level (Extended Data Fig. 6). A per-metric breakdown (Fig. 3b, bottom) shows that SE led across nearly all statistical and biological metrics, with the remaining three models showing more modest differences among themselves in absolute values despite clear rank separation. These results show that the amount of recoverable gene-level signal varies substantially across foundation model embeddings, with SE retaining the most and the remaining three models showing smaller differences.

We then assessed how decoder architecture affects reconstruction quality, aggregating across all four foundation model embeddings (Fig. 3c). For each decoder architecture, we conducted the same pre-registered hyperparameter search as in end-to-end experiments (Supplementary Table 1); for KNN decoders, sensitivity to the number of neighbors showed modest performance changes (Extended Data Fig. 7), so we adopted k = 10 for subsequent analyses. MLP decoders yielded the highest overall scores (Fig. 3c, top), followed by KNN and Transformer decoders. MLP and KNN achieved comparable fidelity on most metrics, whereas Transformer decoders underperformed particularly on biological measures (Fig. 3c, bottom).

An inherent complication in foundation-model reconstruction benchmarking is that pretrained embeddings differ in dimensionality, which can confound reconstruction by changing the information bottleneck available to the decoder. To contextualize these results, we compared the reconstruction performance against the end-to-end PCA baseline at matched dimensionality (Fig. 3d, Extended Data Fig. 8). At their native dimensionalities, foundation models showed mixed performance relative to PCA. SCimilarity at d = 128 outperformed PCA-128 (0.49 vs. 0.40), and scConcept at d = 512 was on par with PCA-512 (0.58 vs. 0.59). In contrast, SE at d = 2,058 and scGPT at d = 512 modestly underperformed PCA at matched dimensionality.

Together, these results show that SE paired with an MLP decoder provides the strongest reconstruction among foundation model–decoder combinations. Decodable signal varies across pretraining objectives and architectures, but is broadly recoverable when paired with a trained decoder. At matched dimensionality, only SCimilarity outperformed PCA, while scConcept was on par and SE and scGPT modestly underperformed, showing that pretraining can but does not uniformly produce embeddings with stronger decodable signal than linear projections.

### High-dimensional PCA performs comparably to foundation model embeddings in latent shift reconstruction

While stand-alone reconstruction performance provides an upper bound on what can be decoded from a given representation, practical applications require latent spaces that remain decodable after manipulation by a downstream model. We therefore assessed reconstruction quality in combination with latent perturbation modeling. Briefly, source cells are encoded into a latent representation via a fixed pretrained encoder, transformed by a perturbation prediction model, decoded back to gene expression via a fixed decoder, and evaluated against true perturbed cells (Fig. 4a).The benchmark primarily compares latent representations, including all end-to-end (PCA, AE, VAE at five dimensionalities) and foundation-model settings (SE embedding from STATE, scConcept, scGPT, SCimilarity), across two perturbation modeling paradigms (STATE and CellFlow), while keeping the representation-specific decoder fixed (see Methods). To match the conventions of perturbation prediction models, which typically operate without explicit library-size handling, we used AE and VAE variants without library-size input for this comparison. To match the use cases in STATE and CellFlow, we evaluated on PBMC-10M using a condition-level split holding out cell type × donor × cytokine combinations (Methods).

**Fig. 4.**
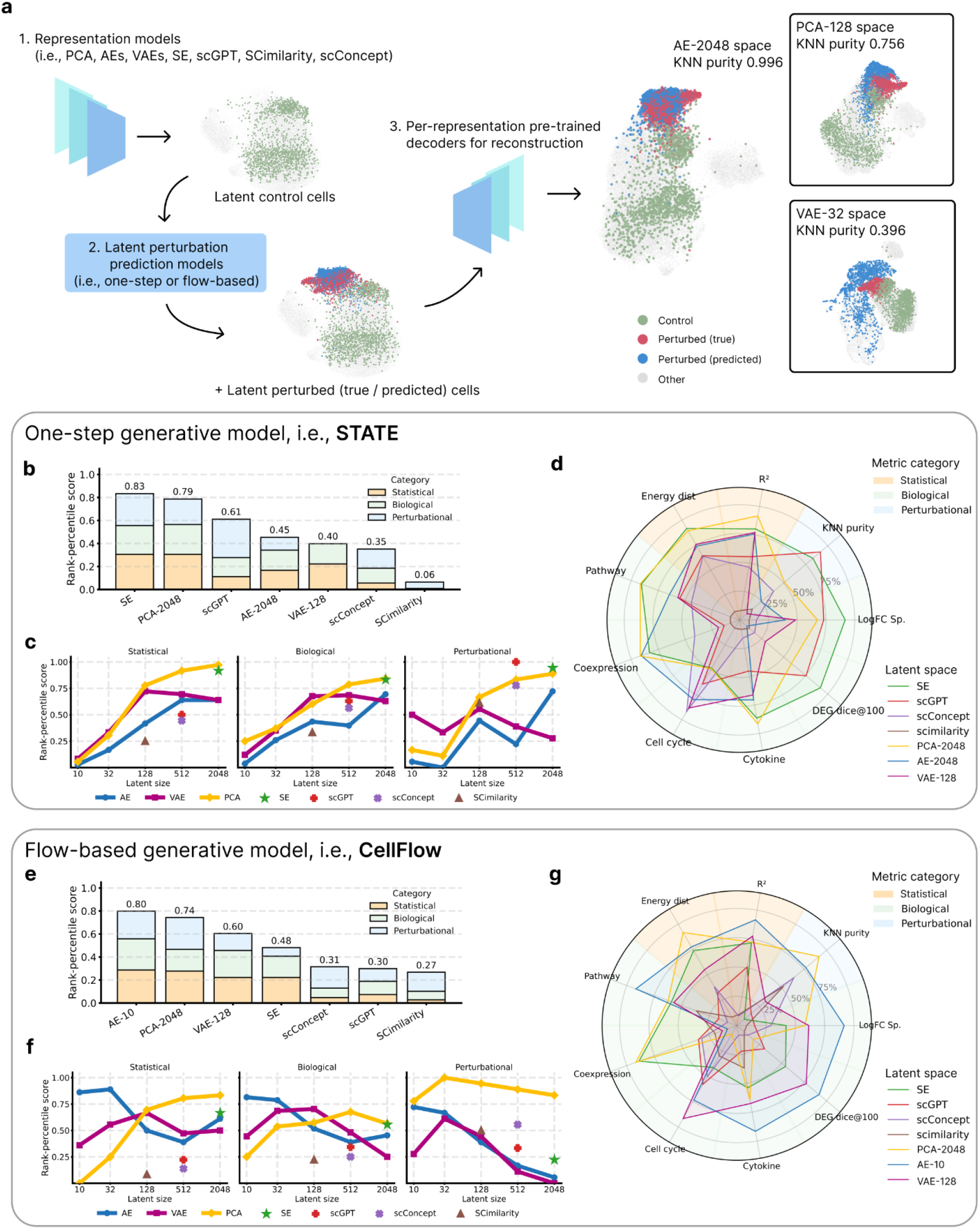
Optimal latent representations for perturbation prediction depend on the downstream model in latent shift reconstruction. **a**, Evaluation pipeline. Source cells are encoded using fixed pretrained encoders, passed through STATE or CellFlow and decoded to gene expression via the corresponding fixed decoder. UMAP visualizations show control, true perturbed, and predicted perturbed cells for representative representation spaces (AE-2048, PCA-128, VAE-32), with corresponding KNN purity. CD14+ monocytes responding to *IFN-β* are shown as the example cell type and perturbation. **b–d**, Reconstruction performance under STATE. **b**, Stacked rank-percentile scores by metric category (statistical, biological, perturbational) for the best-performing latent dimensionality within each end-to-end model family (PCA-2048, AE-2048, VAE-128) and for each foundation model embedding. Bar segments show each category’s contribution to the overall score (computed as the equal-weighted average across categories), with overall scores annotated above each bar. **c**, Rank-percentile score as a function of latent dimensionality for each end-to-end model family (lines), with foundation models (markers) plotted at their native embedding dimensionalities, shown separately for statistical, biological, and perturbational measures. **d**, Per-metric performance shown as a radar plot for the same set of representations. For each metric, values are min-max normalized across the displayed models to [0, 1] with larger radial values representing better performance. **e–g**, Reconstruction performance under CellFlow, displayed as in **b–d**. Best-performing latent dimensionalities are AE-10, PCA-2048, and VAE-128.

Representative UMAP visualizations (Fig. 4a) illustrate the range of reconstruction quality across latent spaces: predicted cells from AE-2048 with STATE overlapped tightly with true perturbed cells (KNN purity = 0.996), while PCA-128 and VAE-32 showed substantially weaker overlap (0.756 and 0.396, respectively). Across both perturbation models, PCA at high dimensionality emerged as a competitive representation. Under STATE, PCA-2048 ranked second only to SE, and under CellFlow, PCA-2048 ranked second only to AE-10, outperforming all four foundation model embeddings (Fig. 4b, e). At the per-metric level, the picture was more nuanced: PCA-2048 led on most metrics in both settings (Fig. 4d, g), while foundation models contributed complementary signals specifically on metrics rewarding perturbation-specific discrimination (i.e., DEG dice@100, LogFC correlation, and KNN purity) under STATE (Fig. 4d), and DEG dice@100 and cell-cycle composition under CellFlow (Fig. 4g). This aligns with our standalone foundation-model reconstruction findings (Extended Data Fig. 8), where foundation models showed relatively strong performance on DEG dice@100. In contrast, VAEs underperformed both PCA and AE families across most metric categories under both perturbation models, with a non-monotonic dependence on latent dimensionality, peaking at d=128 (Fig. 4c, f). Latent space choice effectively affects model performance in reconstructed space (Extended Data Fig. 9, Extended Data Fig. 10) for both models.

### Low-dimensional AE embeddings are optimal for flow-based generative prediction in latent shift reconstruction

The two perturbation models imposed substantially different requirements on representation dimensionality. Under STATE, reconstruction quality improved nearly monotonically with latent dimensionality for end-to-end model families, with the exception of VAE, which peaked at d = 128 (Fig. 4c). Under CellFlow, the dimensionality relationship reversed: AE at d = 10 led on most metrics (Fig. 4e), and increasing AE dimensionality progressively degraded reconstruction quality (Fig. 4f). This trend contrasts with end-to-end reconstruction, where AE performance improved with dimensionality (Fig. 2b). While AE-10 led overall under CellFlow, PCA-2048 retained the strongest performance on the perturbational category specifically (Fig. 4f, right). All foundation model embeddings underperformed end-to-end representations under CellFlow, in contrast to STATE where SE was competitive despite using identical pretrained embeddings in both settings.

Together, these results show that latent space choice is a consequential decision in perturbation prediction. The optimal choice differed substantially between one-step and flow-based generative frameworks: SE and PCA-2048 for STATE, AE-10 and PCA-2048 for CellFlow. No single representation was consistently optimal across both perturbation models, and reconstruction after latent shift diverged from stand-alone reconstruction quality.

## Discussion

We benchmarked reconstruction quality across widely used single-cell representation models and pretrained foundation model embeddings, evaluating end-to-end reconstruction, foundation-model reconstruction, and latent shift reconstruction. Across three datasets, multiple OOD splits, and two perturbation modeling paradigms, our results show that the choice of representation, decoder, and downstream model are interdependent, and that principled selection along all three axes is necessary for faithful gene expression reconstruction from single-cell latent representations.

The strong performance of AEs in stand-alone reconstruction is partly expected, since their objective directly optimizes reconstruction fidelity. However, this observation alone does not trivialize the benchmark, where we address a less obvious question: whether more sophisticated objectives deliver compensating advantages. The regularization imposed by the variational prior, often motivated by its expected benefit for more generalizable embeddings, did not yield more generalizable reconstruction performance over unregularized AEs; in contrast, AEs provided robust reconstruction even under the most out-of-distribution setting. This is notable given that VAE-based representations are widely adopted as the default latent space for perturbation prediction^8,10^. Even PCA, which provides an optimal linear reconstruction baseline, proved competitive at higher dimensionalities and approached AE performance, showing that nonlinear models are not strictly necessary when given sufficient capacity. The same pattern held under latent shift modeling: VAEs underperformed both PCA and AE families across most metric categories under both perturbation models, contradicting the assumption that variational regularization confers an advantage in generative perturbation prediction. These results suggest that, for reconstruction, preserving gene-level structure faithfully matters more than learning smooth or regularized latent geometries, and prompt a reconsideration of the default choice of VAE-based representations when decoding back to gene expression is required.

Among foundation model embeddings, recoverable gene-level information varied with both pretraining objective and embedding dimensionality. SE, which combines the highest dimensionality with a dual-axis loss that explicitly predicts continuous expression values, led overall. Meanwhile, the comparable performance of scConcept and scGPT at matched dimensionality suggests that contrastive cell-level objectives may produce embeddings with decodable signal comparable to gene-level prediction. Decoder architecture also mattered, with MLPs outperforming KNN and Transformer decoders, with Transformers likely suffering from overfitting under limited training data and the high sparsity of scRNA-seq. This is consistent with growing evidence that complex transformer-based architectures do not reliably outperform simpler models on single-cell tasks^37^. The relationship between PCA and foundation model embeddings was nuanced across both foundation-model reconstruction and latent shift reconstruction. In foundation-model reconstruction, SCimilarity surpassed PCA at matched dimensionality, though not all foundation models outperformed PCA. In latent shift modeling, PCA at high dimensionality approached foundation models on most aspects of reconstruction, with foundation models contributing complementary signals on perturbation-separation metrics. The two representations are therefore complementary rather than ranked. Tailoring pretraining objectives to perturbation prediction is a promising direction for future representation learning, with potential to fully leverage foundation models.

In latent shift reconstruction, latent dimensionality emerged as a primary factor governing reconstruction quality, with optimal values depending on the downstream model. STATE leveraged high-dimensional representations, while CellFlow exhibited the opposite pattern for AEs, where low-dimensional embeddings outperformed their high-dimensional counterparts despite weaker stand-alone reconstruction. This likely reflects the distinct mechanisms by which each model class uses the latent space: flow-matching objectives (such as CellFlow’s) require learning continuous transport paths in latent space, making them sensitive to latent dimensionality and geometric structure, while one-step generative frameworks (such as STATE) aggregate information across cells without explicit transport, making them less sensitive to latent geometry but benefiting from higher-capacity representations. Foundation model embeddings, optimized as stand-alone representations, also underperformed end-to-end representations under CellFlow despite remaining competitive under STATE, consistent with this mechanism. Most importantly, stand-alone reconstruction performance does not predict performance after latent-space manipulation: representations that perform well in isolation are not consistently optimal for perturbation prediction. Together, these results show that reconstruction performance reflects a meaningful interaction between representation, inductive bias, and downstream use, rather than merely the training objective.

We summarize practical starting points for representation selection in single-cell reconstruction pipelines. For end-to-end reconstruction, AEs are the default, even when out-of-distribution generalization is required. Reconstruction from pretrained foundation model embeddings is feasible with a separately trained decoder, where MLPs perform best. For latent shift modeling, high-dimensional PCA serves as a robust default across one-step and flow-based generative perturbation modeling paradigms; low-dimensional AE embeddings are preferable for flow-based generative models specifically. Foundation model embeddings are preferred when perturbational separation is the primary concern. These recommendations should be validated as additional perturbation models and datasets become available.

Our work has several limitations. We focused on reconstruction as a necessary property of representations rather than on the broader suitability of latent spaces, including likelihood calibration, sampling quality, or smoothness under interventions. We also considered only log-normalized expression to match common practice; extending the benchmark to count-based decoders (e.g., negative binomial likelihoods) would clarify whether probabilistic count models confer advantages beyond log-space reconstruction. We focused on HVGs due to computational feasibility. Finally, we evaluated foundation models only in their frozen, general-purpose form; evaluating fine-tuned foundation models would disentangle pretraining scale from dataset adaptation, and training cross-dataset AEs at foundation model scale would test directly whether large-scale pretraining improves reconstruction and latent shift robustness.

ReconEval provides a reconstruction-centric perspective that complements existing evaluations of single-cell embeddings focused on integration, annotation, or perturbation prediction accuracy. As latent-space manipulation becomes central to virtual cell modeling, biologically interpretable decoding is essential, yet has lacked systematic evaluation. By introducing a unified metric suite beyond statistical fidelity, we provide both a practical guide and a standardized evaluation framework for representation selection. Our findings also speak to an ongoing shift in representation learning across the broader machine learning community, where embedding-space objectives such as joint-embedding predictive architectures (JEPA^38^) are increasingly adopted over reconstruction-based pretraining. In this context, reconstruction provides a complementary perspective on representation quality. We further incorporate this benchmark as a task within the Open Problems in Single-Cell Analysis platform^39^, enabling comparable evaluation of emerging representation methods as the field moves toward increasingly expressive latent-space models. By framing reconstruction as a downstream task, ReconEval expands the evaluation of single-cell foundation models beyond representation quality toward their ability to support interpretable gene-level predictions. More broadly, we envision this framework strengthening the evaluation of latent space modeling, ensuring that future virtual cell models can be validated by biological experts and ultimately grounded in biology.

## Methods

### Benchmarking setup

#### Latent dimensionality

Since the latent dimension d is a primary bottleneck for reconstruction and directly determines the capacity available for latent-space shift modeling, we benchmarked d ∈ {10, 32, 128, 512, 2048}, covering common settings in single-cell analysis as well as more extreme ones.

#### Hyperparameter selection

We defined a pre-registered hyperparameter search space (Supplementary Table 1, Supplementary Table 2) and selected the best-performing configuration per method on the validation set, with reconstruction loss (MSE) as the selection criterion for end-to-end and foundation-model reconstruction experiments and energy distance in gene-expression space for latent shift experiments.

#### Tasks

We evaluated three reconstruction tasks across two perspectives. Within stand-alone reconstruction: (i) end-to-end reconstruction trained encoder and decoder jointly, comparing PCA, AE, and VAE families; for AE and VAE we additionally evaluated three library-size handling strategies, yielding seven model configurations total. (ii) Foundation-model reconstruction trained a separate decoder to map fixed embeddings from a frozen pretrained foundation model back to gene expression, evaluating embeddings learned from four foundation models (SE embedding from STATE, scGPT, scConcept, SCimilarity) and three decoder architectures (MLP, KNN, Transformer). Within latent shift reconstruction: (iii) the encoder–decoder pairs from both prior settings were coupled with a perturbation prediction model (STATE or CellFlow) operating in latent space.

### Datasets and preprocessing

We benchmarked reconstruction quality on three scRNA-seq datasets spanning perturbational and observational settings.

#### Tahoe-100M

Tahoe-100M is a large-scale cancer cell line perturbation atlas measuring transcriptional responses to small-molecule treatments. It contains 50 cancer cell lines and 379 drugs tested at three dosages, plus one vehicle control (DMSO), yielding 56,877 unique conditions (cell line × drug × dose) and 89,423,257 cells in total. Following the dataset creators, we exclude a technical replicate plate from all experiments to avoid potential leakage across splits; all reported statistics are computed after this exclusion.

#### PBMC-10M

PBMC-10M is a cytokine perturbation dataset profiling immune responses in peripheral blood mononuclear cells. It contains 12 donors and 17 annotated cell types treated with 90 cytokines plus a PBS control, resulting in 17,030 unique conditions (cell type × donor × cytokine) and 9,697,974 cells.

#### LuCA

LuCA is a human lung cancer atlas integrating multiple NSCLC studies. The dataset comprises 892,296 cells across 33 cell types and 1,030 unique conditions (cell type × dataset × tissue origin).

#### Preprocessing

For all datasets, we performed library-size normalization to a fixed target count of 1e4 followed by log(1 + x) transformation, corresponding to scanpy.pp.normalize_total(adata, target_sum=1e4) and scanpy.pp.log1p(adata). For highly variable gene (HVG) selection: on Tahoe-100M, we performed batch-aware HVG selection following established benchmarking practice: we selected the top 3,000 variable genes per plate, and retained genes present in at least two plates. We further included canonical cell-cycle marker genes to ensure cell-cycle metrics are computable, yielding 6,087 genes in total. For PBMC-10M, we followed the dataset-recommended HVG preprocessing, resulting in 5,412 genes. For LuCA, we used the predefined HVG set provided with the integrated atlas, resulting in 5,983 genes. All models were trained and evaluated on the corresponding dataset-specific gene set.

### Split construction

We define a condition as the minimal unit of modeling and evaluation, instantiated as a combination of covariates such as cell type × treatment × dose. All metrics were computed per condition and aggregated. We constructed three train/validation/test splits per dataset to probe generalization at different levels of OOD difficulty, with Split 1 corresponding to the most challenging OOD scenario and Split 3 to the least. All splits were defined using pre-existing dataset covariates with a train/validation/test ratio of 0.70/0.15/0.15 at the level of the held-out unit (Table 1). OOD difficulty of each dataset is quantified in Extended Data Fig. 1.

### Models

Let **X** ∈ ℝ^*M*^ denote a cell-level gene expression vector over *M* genes. Our goal is to evaluate representation learning methods through their ability to support accurate reconstruction of **X** after compression into a low-dimensional latent **Z** ∈ ℝ^*d*^, with *d* < *M*. We formalized each method with an encoder *g*_*ϕ*_ : ℝ^*M*^ → *P* (ℝ^*M*^) that defines a posterior *q*_*ϕ*_ (**Z** | **X**) over latent variables, and a decoder *f*_*θ*_ : ℝ^*M*^ → *P* (ℝ^*M*^) that defines a likelihood *q*_*θ*_ (**Z** | **X**). Reconstruction quality was assessed by comparing samples from *p*_*θ*_ (**X** | **Z**) (with **Z** ∽*q*_*ϕ*_ (**Z** | **X**)) against the observed data.

### End-to-end reconstruction

#### PCA

PCA serves as a linear baseline, which yields the optimal rank-*d* reconstruction in the least-squares sense. We treat PCA as a reference point for both reconstruction fidelity and the benefits of nonlinear modeling.

#### Autoencoder (AE)

AE parameterizes *g*_*ϕ*_ and *f*_*θ*_ as deterministic deep neural networks trained to minimize a reconstruction objective. The approximate posterior degenerates to a point mass *q*_*ϕ*_ (**Z** | **X**) = *δ(***Z** − *g*_*ϕ*_ (**X**)).

#### Variational autoencoder (VAE)

VAE further defines a stochastic latent variable model with a Normal distribution prior. We adopted scVI as our VAE implementation. The encoder defines a diagonal Gaussian posterior 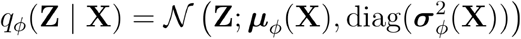, and the model is trained by maximizing the evidence lower bound (ELBO). We also considered a weighted-KL variant (*β* -VAE)^40^.

#### Library-size handling

Sequencing depth (library size) is a major technical factor that can be treated either as a technical covariate or as part of the biological signal, and different modeling choices can substantially affect both reconstruction and downstream use. To isolate this axis, we evaluated three common strategies as in scVI implementation^4^: (i) *None*: no library size information is provided to the model; the decoder must implicitly account for depth variation during reconstruction. (ii) *Modeled*: library size is a learned cell-level variable inferred from X during training. For AEs, we predicted library size using a single linear layer. For VAEs, we introduced an additional latent variable with a data-driven prior and corresponding KL term. (iii) *Observed*: true library size is provided explicitly as an input covariate to the decoder. For PCA, only the *none* strategy was applicable. These strategies reflect common choices in single-cell analysis pipelines, where sequencing depth can be treated as a technical covariate to correct for, a latent variable to infer, or simply left implicit in the representation.

### Foundation-model reconstruction

#### SE (Embeddings learned from STATE)

SE is the encoder component of the STATE framework^17^. The model is a bidirectional transformer, producing a cell embedding of dimension d = 2,058. The loss consists of mean squared error between predicted and true log-normalized expression values, computed both within each cell and across cells within each minibatch. SE is pretrained on 167 million observational human cells.

#### scGPT

scGPT^26^ is a generative pretrained transformer, producing a cell embedding of dimension d = 512. The loss consists of mean squared error between predicted and true binned expression values, combined across two objectives: gene expression prediction (GEP) from unmasked gene context, and gene expression prediction conditioned on the cell embedding (GEPC). scGPT is pretrained on over 33 million cells.

#### Scimilarity

SCimilarity^24^ is a feedforward network, producing a cell embedding of dimension d = 128 constrained to lie on the unit hypersphere. The loss consists of a weighted combination of a supervised triplet loss (using cell-type annotations to embed same-type cells close together) and an auxiliary MSE reconstruction loss. SCimilarity is pretrained on 22.7 million annotated cells.

#### scConcept

scConcept^25^ is a transformer-based contrastive learning framework, producing a cell embedding of dimension d = 512. The loss consists of an InfoNCE-style contrastive objective that pulls together computational views of the same cell while pushing apart views of different cells, without any gene-level reconstruction term. scConcept is pretrained on over 30 million annotated cells.

#### Foundation-model-specific preprocessing

SCimilarity inputs were aligned to the model’s fixed feature space and log-normalized using the provided preprocessing utilities. STATE inputs were log1p-transformed following the official interface. For scGPT and scConcept, inputs were discretized via value binning and/or rank-based encodings as part of their modeling pipelines; we followed the same preprocessing scheme defined in each study.

#### Decoder architectures

Given a fixed embedding 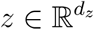 from a frozen foundation model encoder, we trained three decoder architectures to reconstruct gene expression *x* ∈ ℝ^*G*^:

MLP Decoder. A multilayer perceptron with *L* hidden layers of width *H*, batch normalization, and ReLU activations:

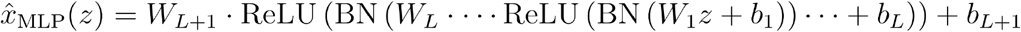

with hidden depth and width swept as in end-to-end decoders (Supplementary Table 2).

KNN Decoder. A non-parametric lookup that retrieves the *k* nearest training cells in embedding space and averages their gene expression:

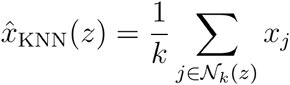

where *N*_*k*_(*z*) is the set of indices of the *k* training cells with smallest Euclidean distance to *z* in embedding space. We used *k* = 10 for the main analyses (sensitivity analysis in Extended Data Fig. 7).

Transformer decoder. A gene-aware transformer that predicts expression for each gene conditioned on the cell embedding:

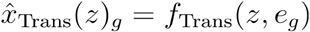

where *e*_*g*_ is a learned gene embedding indexed by gene *g* ∈ {1,…,*G*}, and *f*_Trans_ is a multi-head transformer with *l* layers, *h* attention heads, and inner dimension *d*_hid_ . We compared a small ({4 attention heads, 3 layers}) and large ({8 attention heads, 6 layers}) configuration (Supplementary Table 2). All parametric decoders (MLP, Transformer) were trained from scratch on each dataset split using mean squared error loss against log-normalized gene expression. Hyperparameters were selected by validation loss.

### Latent shift reconstruction

#### STATE

A one-step transformer-based generative model that predicts perturbation effects across cell populations, matching predicted and ground-truth populations by minimizing their energy distance in both the latent space and gene-expression space.

#### CellFlow

A flow-based generative framework that models perturbation-induced phenotypic changes via learned continuous transport paths from unperturbed to perturbed cell states in the latent space.

For each combination of latent space and perturbation model, source cells were encoded using the fixed pretrained encoder, shifted in latent space by the perturbation model, and decoded to gene expression via the corresponding fixed decoder. To align with both models’ native evaluation settings, we evaluated on PBMC-10M using a condition-level split (cell type × donor × cytokine).

### Metrics

We evaluated reconstruction quality using statistical, biological and perturbational metrics. For one run, all metrics were computed per condition (e.g., cell type × drug × dose) and mean aggregated across conditions. Statistical and biological measures were computed for all three tasks; the perturbational measure (KNN purity) was computed only for latent shift reconstruction. Cytokine response correlation was computed only for PBMC-10M and LuCA, where the cytokine reaction context applies. Due to the computational cost of evaluating across the full PBMC-10M and Tahoe-100M test sets (1.5M and 15M cells respectively), all metrics were computed with 5% subsampling. To mitigate sampling variability, evaluation was repeated across 5 independent random seeds, and reported scores are the mean across seeds.

### Statistical measures

Single-cell measurements often exhibit substantial stochasticity and technical noise. Thus, per-cell correspondence between an observed cell and its reconstruction is generally ill-posed. We complemented cell-level with population-level evaluation of the reconstruction output within each condition.

#### MSE

When a deterministic point reconstruction 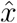 is available, we computed the mean squared error between *x* and 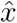:

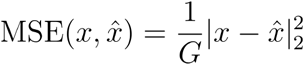

where *G* is the number of genes.

#### R^2^

Let *X* and 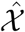 denote the sets of ground-truth and reconstructed cells for a condition, where each cell is a gene expression vector. We computed the coefficient of determination between condition-wise mean expression vectors:

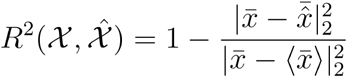

where 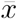 and 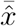 are the condition-wise mean expression vectors of *X* and 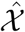, respectively.

#### Energy dist

For higher-order distributional agreement, we first mapped cells into a shared low-dimensional evaluation space using PCA fit on all ground-truth cells, then computed the energy distance in this PCA space:

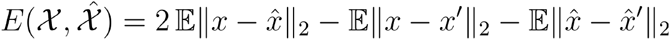

where *x, x*′ ∽ *X*’ are independent samples from the ground-truth cell population, 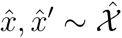 are independent samples from the reconstructed cell population, ║ · ║ denotes the Euclidean norm in PCA space, and expectations are approximated by empirical averages over all pairs.

### Biological measures

We quantified whether reconstructions still preserve biologically meaningful variation through proxies of common downstream analyses. For perturbation-based metrics, we denote ground-truth perturbed cells for a condition as *X*, corresponding ground-truth control cells as *R*, and their reconstructed counterparts as 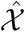 and 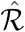.

#### DEG dice@k

For each perturbation condition, we identified differentially expressed genes (DEGs) between perturbed and control populations using the Wilcoxon rank-sum test on log-normalized expression, performed independently on ground-truth and reconstructed data using scanpy.tl.rank_genes_groups. We computed the Dice coefficient between the top-*k* DEGs from ground-truth and reconstructed data, where genes were ranked by absolute log-fold-change after pre-filtering for statistical significance (Benjamini-Hochberg adjusted *p* -value ≤ 0.05):

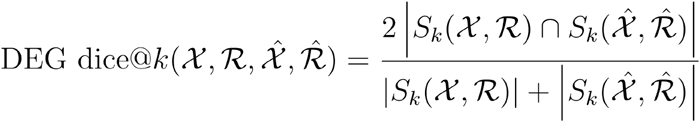

where *S*_*k*_(·, ·) denotes the top *k* genes ranked by absolute log-fold-change among genes passing the FDR threshold from the differential expression contrast between two cell populations, and ⋂ denotes set intersection. We report results at *k* = 100, which we found to provide a stable balance between sensitivity (capturing the most biologically meaningful DEGs) and noise (avoiding the inclusion of marginally significant genes).

#### LogFC Sp

We computed the Spearman rank correlation between log-fold-changes from ground-truth and reconstructed contrasts across all genes with defined values in both:

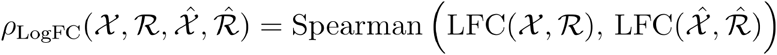

where LFC (·, ·) denotes the gene-wise log-fold-change between two cell populations, computed using scanpy.tl.rank_genes_groups with the same Wilcoxon rank-sum test as for DEG dice. The correlation was computed across all genes with defined log-fold-change values in both contrasts. This complements DEG dice@k by capturing the global rank correspondence of perturbation effects across the full transcriptome rather than overlap restricted to top-ranked genes, providing a transcriptome-wide perspective on whether reconstruction preserves the relative ordering of gene-level effects.

#### Pathway

We computed pathway activity scores using pre-defined responsive genes (PROGENy^41^) via univariate linear model (ULM) scoring on reconstructed and ground-truth cells, and reported Spearman correlation between ground-truth and reconstructed scores, averaged across valid pathways:

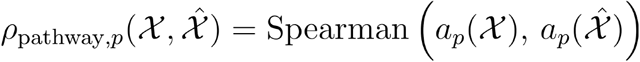

where *a*_*p*_(·) denotes the ULM activity score of pathway *p* computed across cells. The aggregated pathway score across pathways is:

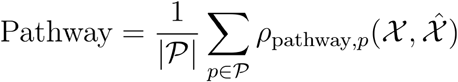

where *P* is the set of valid pathways.

#### Coexpression

We evaluated the preservation of gene-gene correlation structure by computing Spearman coexpression matrices for curated gene sets (MSigDB Hallmark collection^42^) on reconstructed and ground-truth cells, accessed via OmniPath^43^. For each gene set, we computed the per-pair similarity between ground-truth and reconstructed correlations, then averaged across pairs:

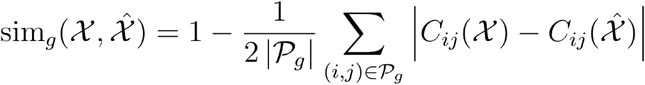

where *C*_*ij*_(·) is the gene-gene Spearman correlation between genes *i* and *j*, and *P*_*g*_ is the set of unordered gene pairs in gene set *g*. The factor of 1/2 normalizes the absolute difference (which ranges over [0, 2] since correlations lie in [-1, 1]) to a similarity score in [0, 1]. The aggregated coexpression score across gene sets is:

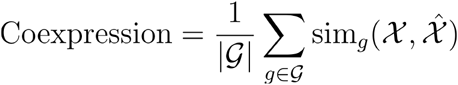

where *g* is the collection of gene sets.

#### Cytokine

We assessed cytokine response preservation by computing cytokine activity scores from cytokine-specific gene signatures from the Immune Dictionary^44^ using scanpy.tl.score_genes. For each cell population, we computed one score per cytokine as the mean of cell-level scores. For each cell type, we then computed the Spearman correlation between true and reconstructed cytokine score vectors across cytokines applicable to that cell type:

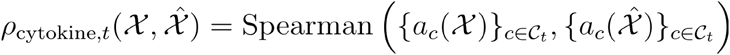

where *a*_*c*_(·) is the mean cytokine activity score for cytokine *c* in a population, and *C*_*t*_ is the set of cytokines applicable to cell type *t* in the Immune Dictionary. The aggregated cytokine score across cell types is:

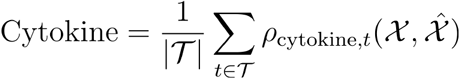

where *T* is the set of cell types with applicable cytokine signatures.

#### Cell cycle

We assessed whether cell-cycle composition is preserved by computing canonical cell-cycle scores from established marker genes^33^ using scanpy.tl.score_genes_cell_cycle, which scores each cell for S-phase and G2/M-phase gene expression and assigns a phase label (G1, S, or G2/M). We reported the proportion of cells retaining their phase assignment after reconstruction:

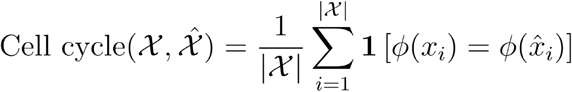

where *ϕ* (·) is the phase assignment from canonical marker gene scoring, *x*_*i*_ and 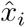 are paired ground-truth and reconstructed cells, and 1[·] is the indicator function returning 1 if the condition holds and 0 otherwise.

### Perturbational measure

#### KNN purity

We assessed whether predicted perturbed cells retain perturbation-specific effects in gene expression space. We defined a KNN purity score that measures the overlap between predicted cells and true perturbed cells relative to control cells. Let *R* = *X*^*ctrl*^ ∪ *X*^*pert*^ denote the reference pool composed of true control and true perturbed cells. For a query set *Q*, we computed the fraction of its *k*-nearest neighbors in that *R* belong to the true perturbed population:

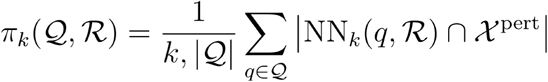

We then normalized this score relative to a control baseline and an upper bound derived from the true perturbed cells themselves:

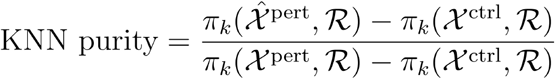

A score of 0 indicates that predicted cells are indistinguishable from control cells (no perturbation effect retained), while a score of 1 indicates that predicted cells overlap with true perturbed cells to the same degree as true perturbed cells overlap with themselves. We used *k* = 10 and computed the metric in gene expression space.

### Rank-percentile score

To provide a robust summary of overall performance across heterogeneous metrics, we defined a rank-based aggregate score. For each (dataset, split, latent dimensionality) combination, we ranked methods per metric, with MSE and energy distance inverted so that higher values indicate better performance throughout. Ranks were converted to rank-percentile scores rp_*i*_(*m*) in [0,1] as:

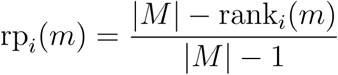

where rank_*i*_(*m*) denotes the rank of method *m* on metric *i* (rank 1 = best) and |*M*| is the number of methods.

Rank-percentile scores were first averaged within each metric category (statistical, biological, perturbational), then averaged across categories to obtain the overall rank-percentile score. This two-stage aggregation prevents categories with more metrics from dominating the overall score.

### Software and reproducibility

All preprocessing was performed using scanpy^45^. VAE models were implemented using scVI^4^. Pathway activity scores were computed using decoupler with PROGENy^41^.

Coexpression gene sets were obtained from MSigDB (Hallmark collection)^42^ accessed via OmniPath^43^. Cell-cycle scoring used marker genes from Tirosh et al 2016^33^.

## Data availability

The three datasets used in this study are publicly available from their original sources: Tahoe-100M^34^ at https://huggingface.co/datasets/tahoebio/Tahoe-100M, the Human Cytokine Dictionary (PBMC-10M)^35^ at https://www.parsebiosciences.com/datasets/10-million-human-pbmcs-in-a-single-experiment/, and the Single-Cell Lung Cancer Atlas (LuCA)^36^ via CellxGene Discover (version: 240701). Preprocessing scripts, including highly variable gene selection and train/validation/test split definitions used in our benchmark, are available alongside the code repository at https://github.com/theislab/ReconEval.

## Code availability

The benchmark code, including all preprocessing pipelines, reconstruction models, metric implementations, and analysis notebooks, is publicly available at https://github.com/theislab/ReconEval.

## Acknowledgements

We thank S. Becker for designing the cytokine metric, L. Heumos and W. Wang for reviewing the paper, F. Fischer and A. A. Monifar for valuable feedback on project design, L. Sikkema for introducing us to the LuCA dataset, I. Gold for suggestions on the Zarr implementation enabling scVI scaling to Tahoe-100M, M. Minaeva for feedback on figures, and T. Richter for help with interpreting the PCA results. This work was supported by the Deutsche Forschungsgemeinschaft (DFG) - Project number 513025799 (grant number TH 900/19-1). Funded by the European Union (ERC, DeepCell - 101054957). This project has received funding from the European Research Council (ERC) under the European Union’s Horizon Europe research and innovation programme under grant agreement No. 101248740 (CellCourier). This work was supported by the Helmholtz Association within the framework of the Helmholtz Foundation Model Initiative (VirtualCell). This work was funded by the Bavarian State Ministry of Economic Affairs, Regional Development and Energy under the Bavarian Collaborative Research Program (BayVFP) for the period 2024–2027. This project has received funding from the Free State of Bavaria’s Hightech Agenda through the Institute of AI for Health (AIH). This work was supported by the Chan Zuckerberg Initiative Foundation (CZIF; grant CZIF2022-007488 (Human Cell Atlas Data Ecosystem)).

## Author contributions

X.F., D.K., and A.P. wrote the paper. D.K., X.F., A.P., and A.T.L. designed the study. X.F., D.K., A.P., and A.T.L. designed the evaluation. X.F. and E.A. prepared the experiments. X.F., D.K., E.A., and A.P. prepared the figures. X.F. and E.A. implemented the metrics and wrote the code. M.B. implemented the transformer decoder. L.B.K. implemented the coexpression metric. M. L. helped with the Tahoe-100M analysis. D.K., A.P., M.D.L., and F.J.T. supervised the work. All authors reviewed the final paper.

## Extended Data

**Extended Data Fig. 1.**
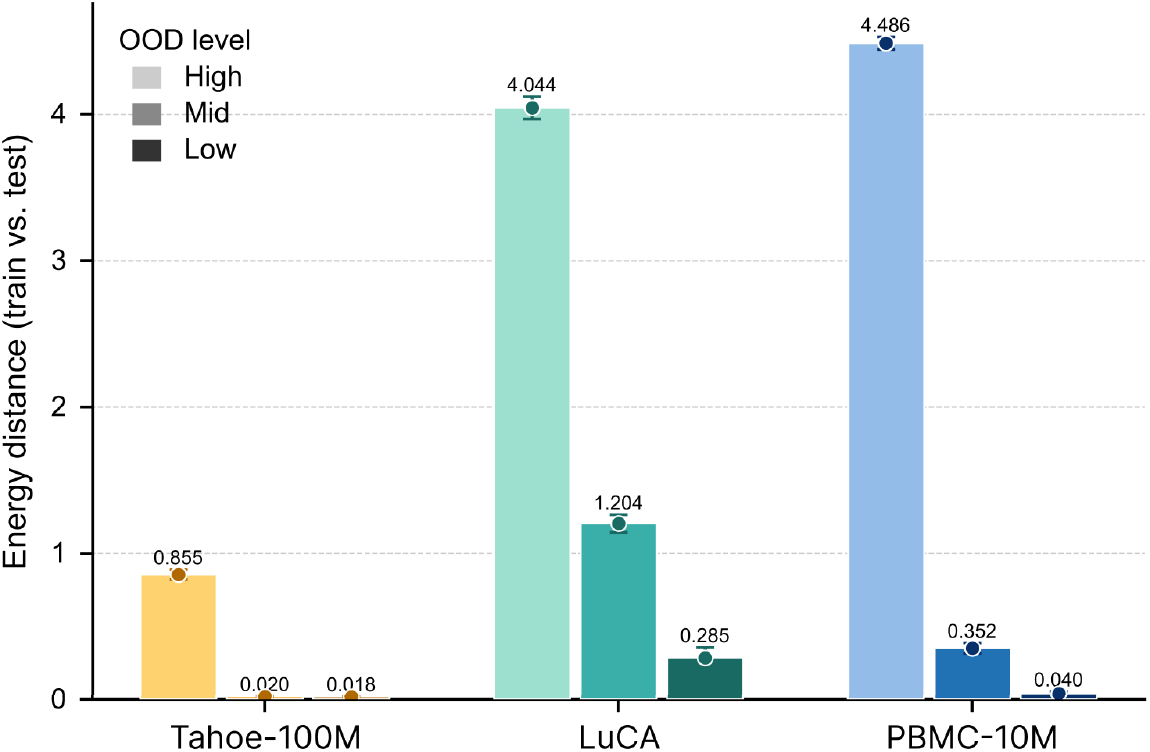
Out-of-distribution (OOD) levels across train–test splits for each benchmark dataset. Energy distance between train and test sets in log-normalized expression space, computed on the top 50 principal components, for the three benchmark datasets across three OOD difficulty levels (high, mid, low). For Tahoe-100M, splits hold out cell lines (high), drugs (mid), or cell line × drug × dose combinations (low). For LuCA, splits hold out cell types (high), datasets (mid), or cell type × dataset × tissue combinations (low). For PBMC-10M, splits hold out cytokines (high), donors (mid), or cell type × donor × cytokine combinations (low). Higher energy distance indicates greater distributional shift between train and test data. Bars show mean ± standard error across 5 random subsamples of 1,000 cells each.

**Extended Data Fig. 2.**
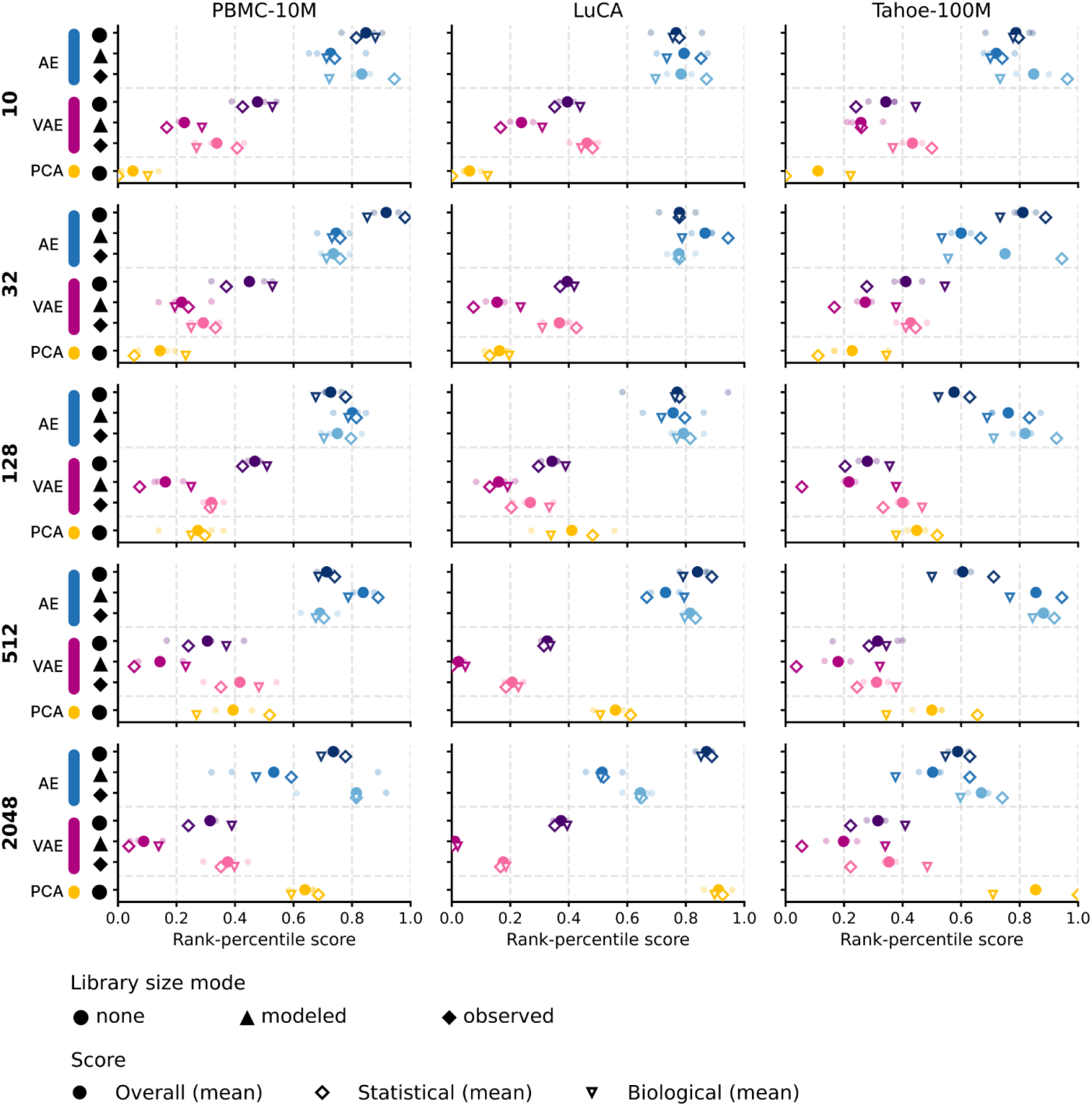
End-to-end reconstruction performance across datasets and latent dimensionalities. Rank-percentile scores for end-to-end reconstruction methods (AEs, VAEs, PCA) on Tahoe-100M, PBMC-10M, and LuCA (columns), evaluated at five latent dimensionalities d = 10, 32, 128, 512, 2,048 (rows). For each model X library-size combination, we report rank-percentile scores for statistical fidelity (◇) and biological signal preservation (▽); the overall score (◆) is their equally weighted average. Solid markers denote the mean across OOD splits, while transparent dots show individual split values. Library-size handling (none, modeled, observed) is indicated by marker offset within each row. Results are aggregated across all OOD splits per dataset; higher rank-percentile score indicates better performance.

**Extended Data Fig. 3.**
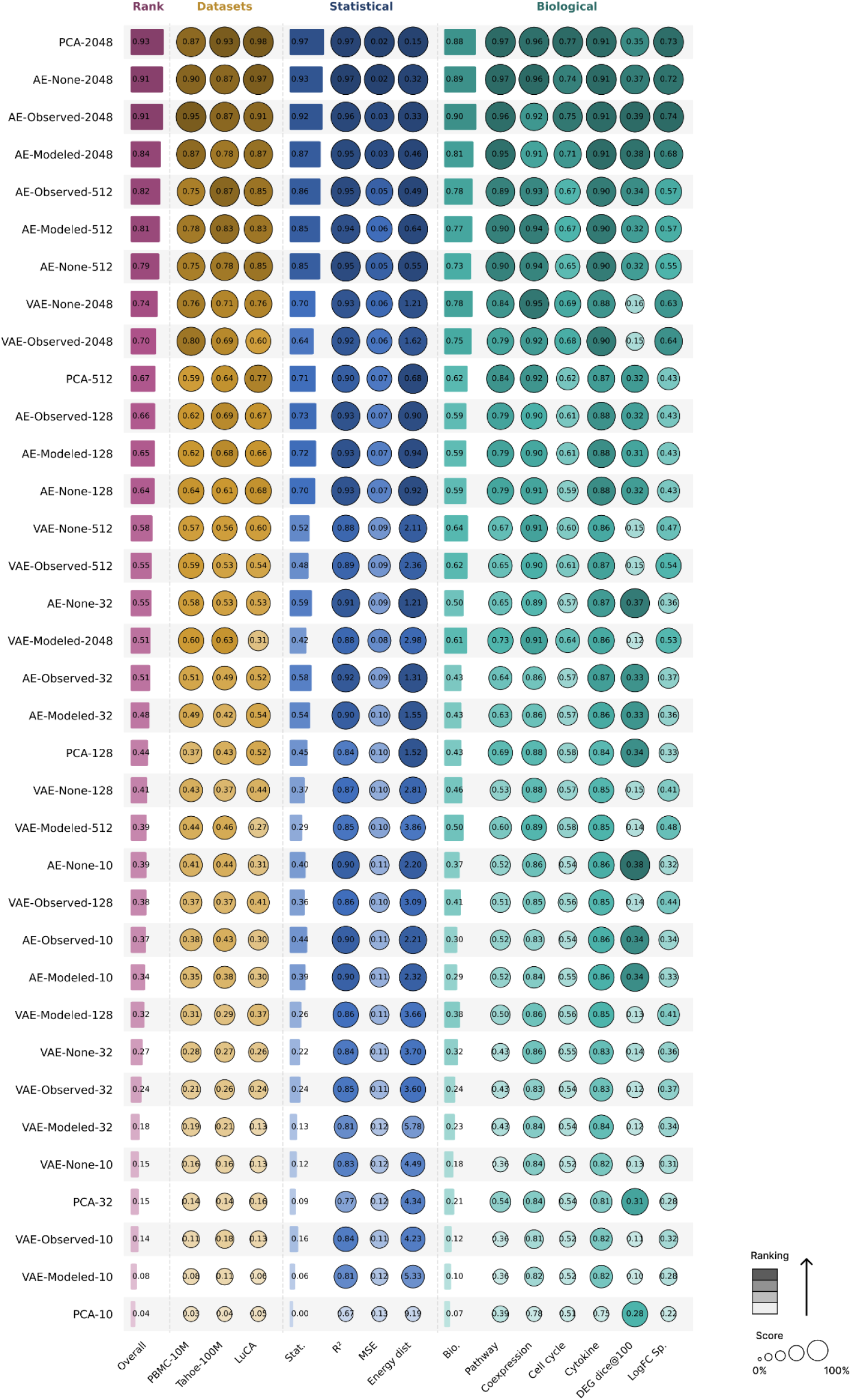
End-to-end reconstruction across architectures, dimensionalities, and library-size handling strategies. Reconstruction performance on PBMC-10M, Tahoe-100M, and LuCA for PCA, AE, and VAE at five latent dimensionalities (d = 10, 32, 128, 512, 2,048), aggregated across OOD splits. AE and VAE rows are labeled with library-size mode (None, Modeled, Observed; see Methods). Higher values are better for all metrics except MSE and Energy distance.

**Extended Data Fig. 4.**
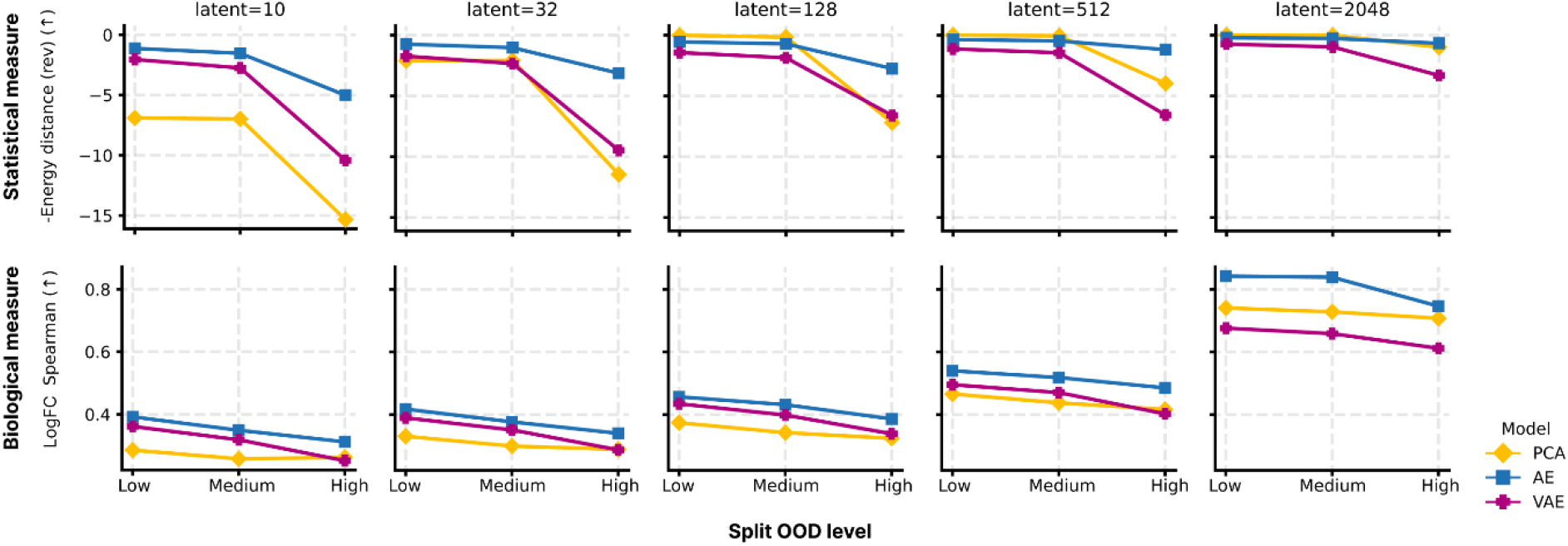
Robustness of end-to-end reconstruction to increasing OOD difficulty on PBMC-10M. PBMC-10M was selected as the example dataset because it shows the largest separation between OOD levels (Extended Data Fig. 1). Reconstruction performance for AE, VAE, and PCA on PBMC-10M across three out-of-distribution difficulty levels (low, medium, high; Split 3, Split 2, Split 1 in Table 1) and five latent dimensionalities. Top: negative energy distance (an example statistical measure), where higher values indicate closer match between reconstructed and true expression distributions. Bottom: LogFC Spearman correlation across differentially expressed genes (an example biological measure), where higher values indicate better recovery of perturbation-induced expression changes. Performance generally degrades with increasing OOD difficulty across all model families, though the magnitude of degradation varies with latent dimensionality.

**Extended Data Fig. 5.**
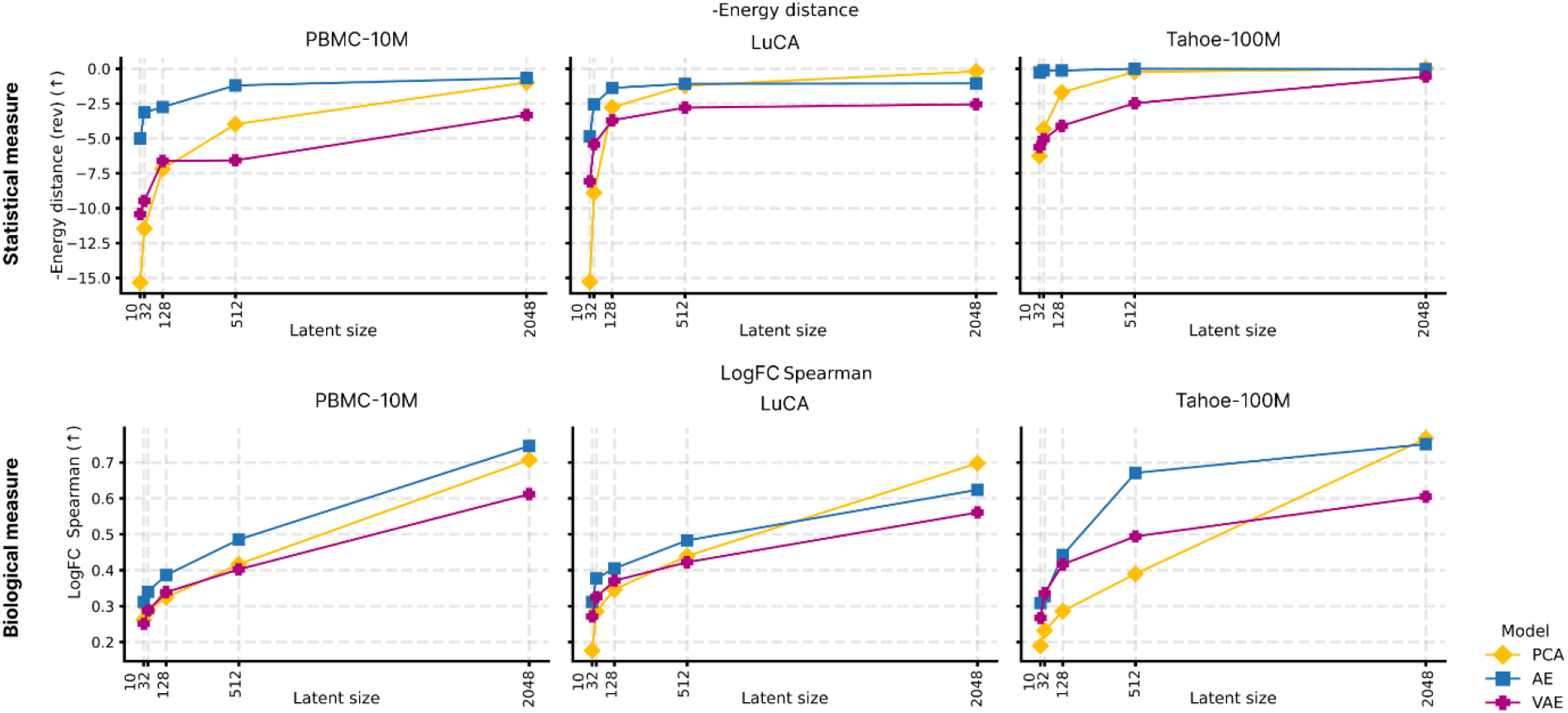
Scaling of end-to-end reconstruction performance with latent dimensionality across datasets. Reconstruction performance for AE, VAE, and PCA across five latent dimensionalities on PBMC-10M, LuCA, and Tahoe-100M (columns), evaluated on the most challenging OOD split (Split 1, holding out cell types or cell lines). Top: negative energy distance (an example statistical measure), where higher values indicate closer match between reconstructed and true expression distributions. Bottom: LogFC Spearman correlation across differentially expressed genes (an example biological measure), where higher values indicate better recovery of perturbation-induced expression changes. For each model family, the best-performing library-size handling strategy is shown (modeled for AE, none for VAE).

**Extended Data Fig. 6.**
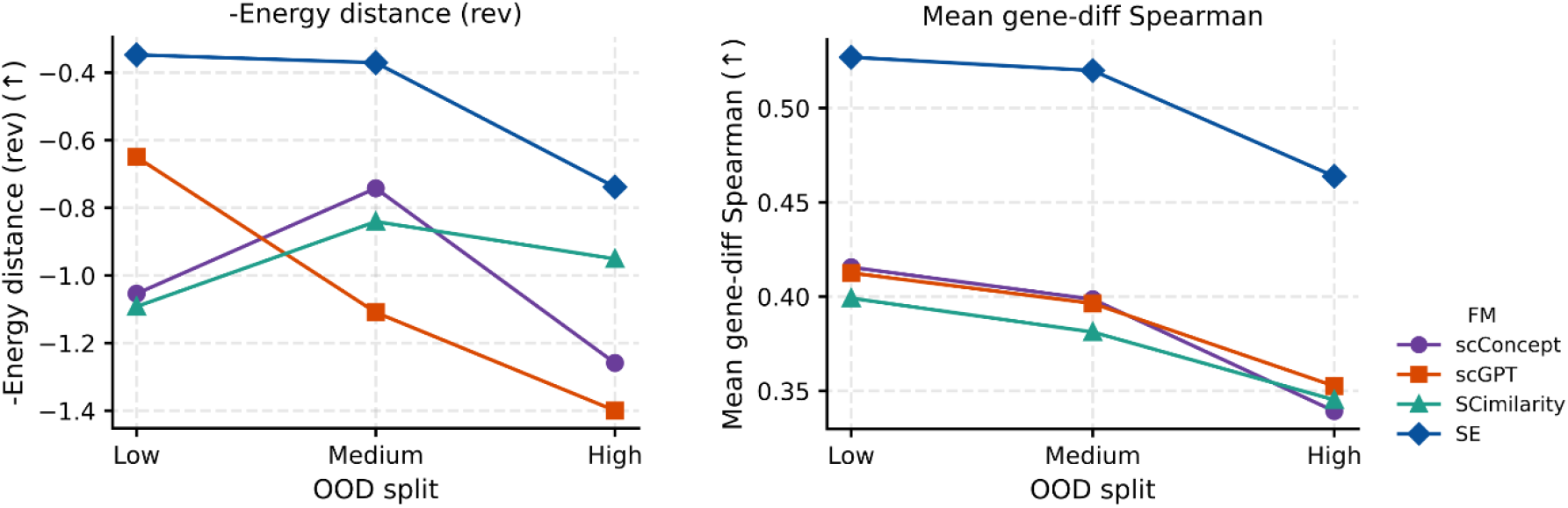
Out-of-distribution robustness of foundation model embeddings under foundation-model reconstruction on PBMC-10M. Reconstruction performance as a function of OOD split difficulty (low / medium / high) for four foundation model embeddings (SE, scConcept, scGPT, SCimilarity) paired with MLP decoders, illustrated by Energy distance (statistical, reversed to be larger better) and LogFC Spearman correlation (biological).

**Extended Data Fig. 7.**
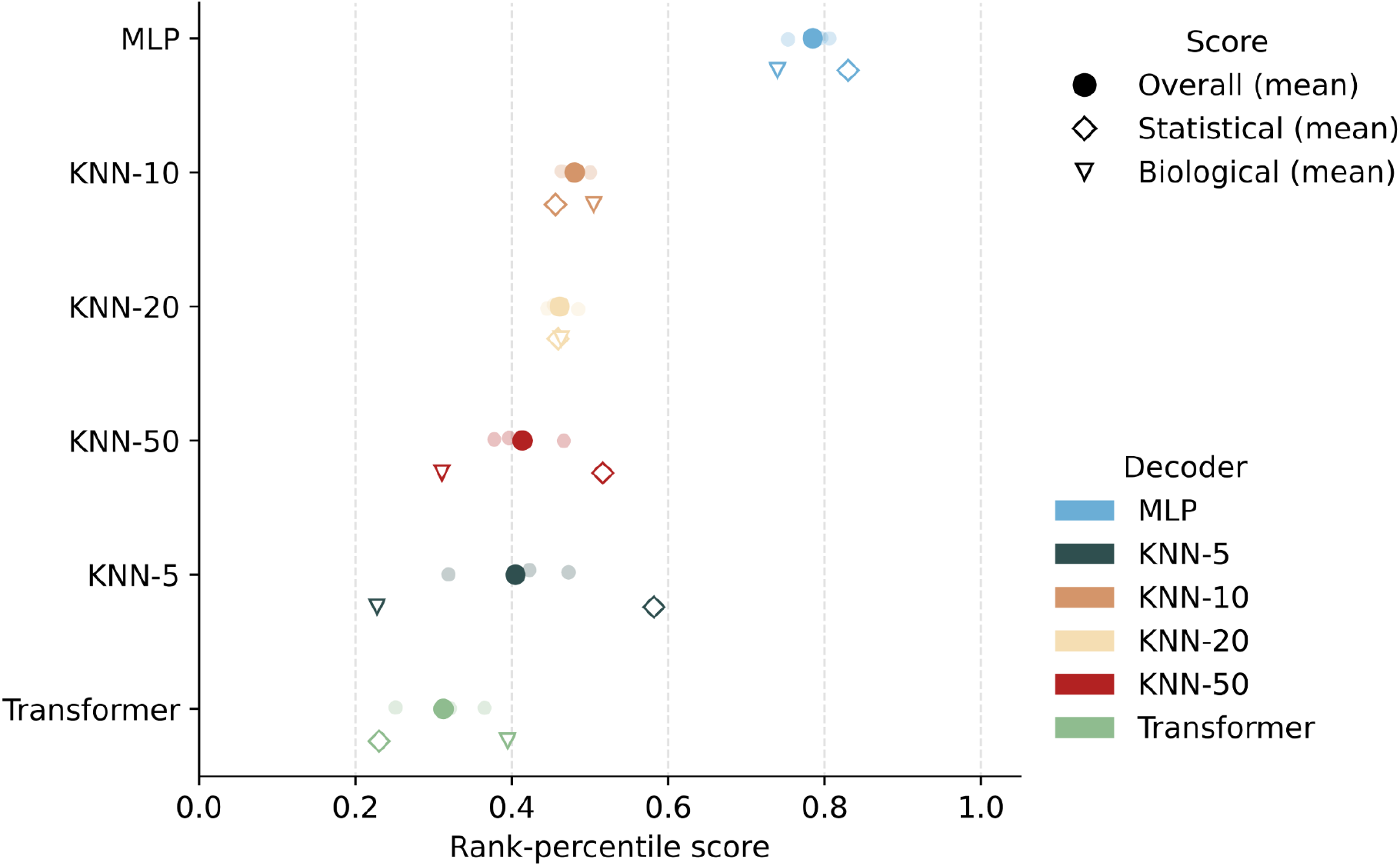
Decoder architecture and KNN neighbor sensitivity for scConcept embeddings. Rank-percentile scores for reconstruction from scConcept embeddings on PBMC-10M, comparing MLP, KNN (at k = 5, 10, 20, 50 neighbors), and Transformer decoders. Solid markers denote the mean across OOD splits, while transparent dots show individual split values. For each decoder, we report rank-percentile scores for statistical fidelity (◇) and biological signal preservation (▽); the overall score (◆) is their equally weighted average. Relative performance of KNN varied modestly across k∈{5,10,20,50}. MLP decoders consistently outperformed both KNN and Transformer decoders across score categories.

**Extended Data Fig. 8.**
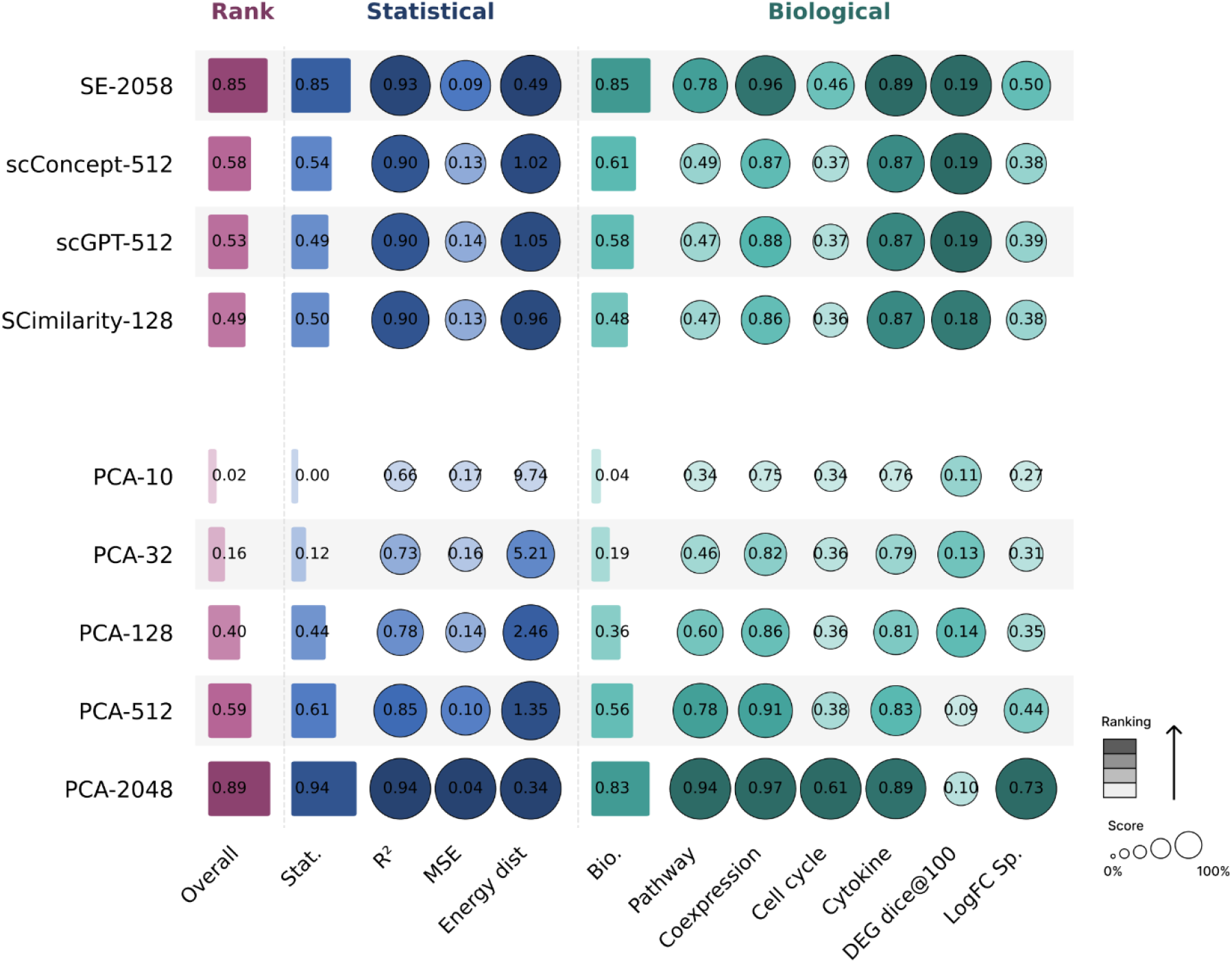
Foundation-model reconstruction of foundation model embeddings compared to end-to-end PCA. Reconstruction performance on PBMC-10M for four foundation models with MLP decoders (SE, scConcept, scGPT, SCimilarity) and PCA at five latent dimensionalities (d = 10, 32, 128, 512, 2048), aggregated across out-of-distribution (OOD) splits. The Rank column shows aggregate rank-percentile, broken down into statistical and biological subcategories. Higher values indicate better performance for all metrics except MSE and energy distance, where lower is better. Bar lengths and circle sizes both scale with score magnitude.

**Extended Data Fig. 9.**
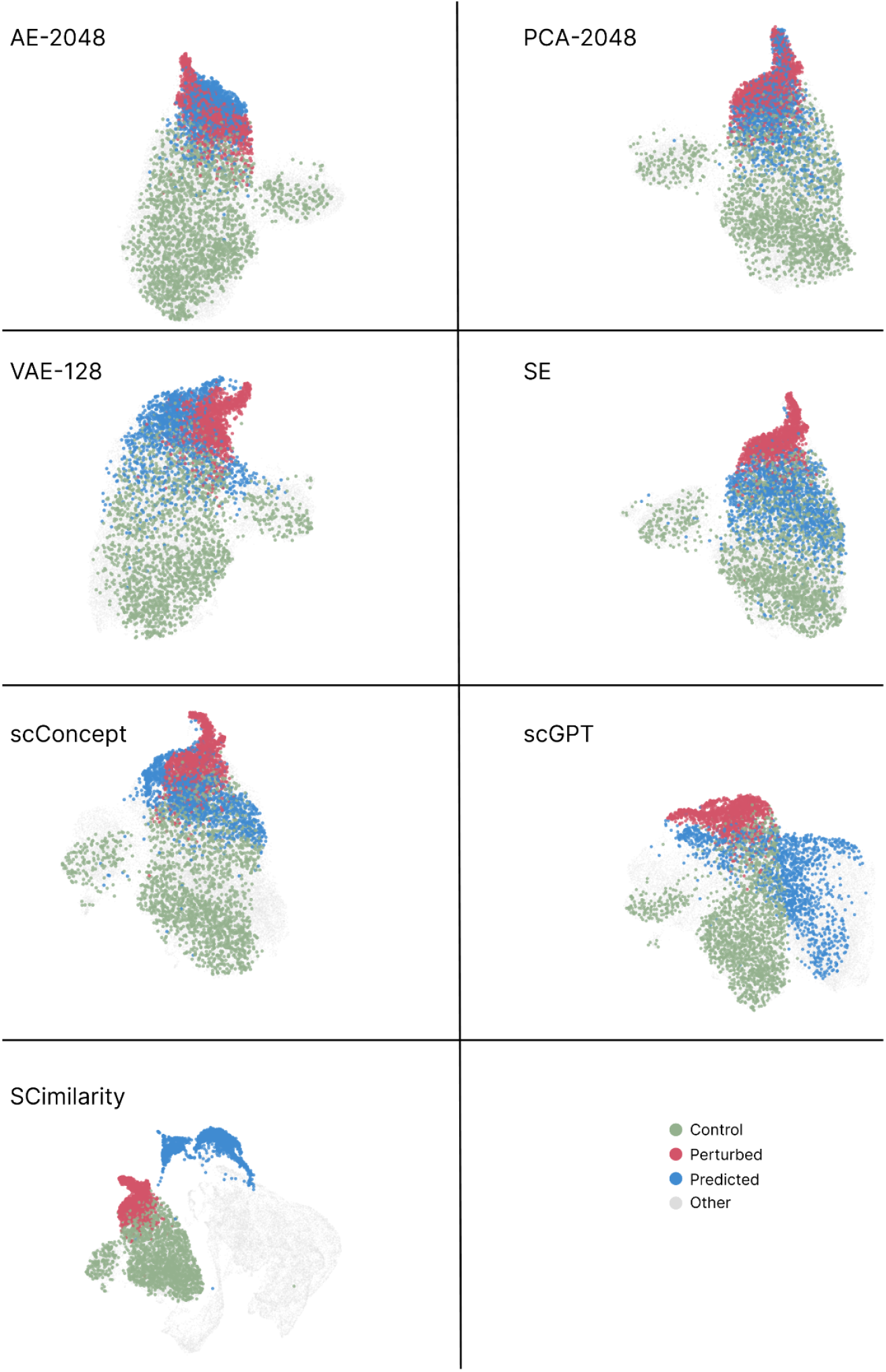
Qualitative comparison of latent shift reconstruction with STATE across representations. UMAP projections of true control cells, true perturbed cells, predicted perturbed cells, and other cells from the dataset (gray) for the *IFN-β* perturbation on CD14+ monocytes from PBMC-10M. Seven latent representations are shown: end-to-end models at their best-performing dimensionality under STATE (AE-2048, PCA-2048, VAE-128) and four foundation model embeddings (SE, scConcept, scGPT, SCimilarity). UMAPs were computed in a 50-dimensional PCA space fit on the combined set of true control, true perturbed, and predicted cells (downsampled for visualization). Closer overlap of predicted cells with true perturbed cells indicates better reconstruction of perturbation effects in gene expression space.

**Extended Data Fig. 10.**
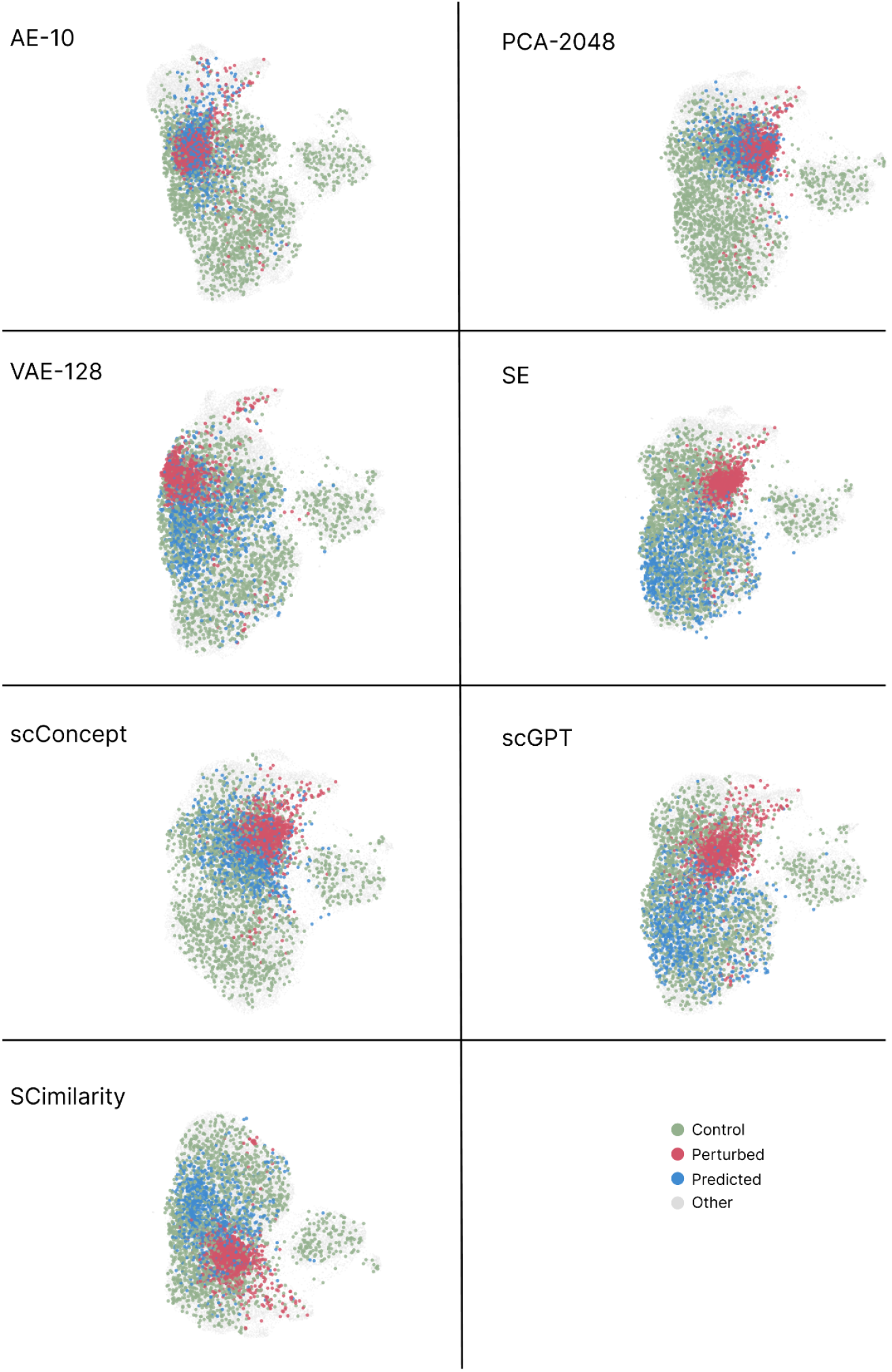
Qualitative comparison of latent shift reconstruction with CellFlow across representations. UMAP projections of true control cells, true perturbed cells, predicted perturbed cells, and other cells from the dataset (gray) for the *IFN-γ* perturbation on CD14+ monocytes from PBMC-10M. Seven latent representations are shown: end-to-end models at their best-performing dimensionality under CellFlow (AE-10, PCA-2048, VAE-128) and four foundation model embeddings (SE, scConcept, scGPT, SCimilarity). UMAPs were computed in a 50-dimensional PCA space fit on the combined set of true control, true perturbed, and predicted cells (downsampled for visualization). Closer overlap of predicted cells with true perturbed cells indicates better reconstruction of perturbation effects in gene expression space.

**Supplementary Table 1.**
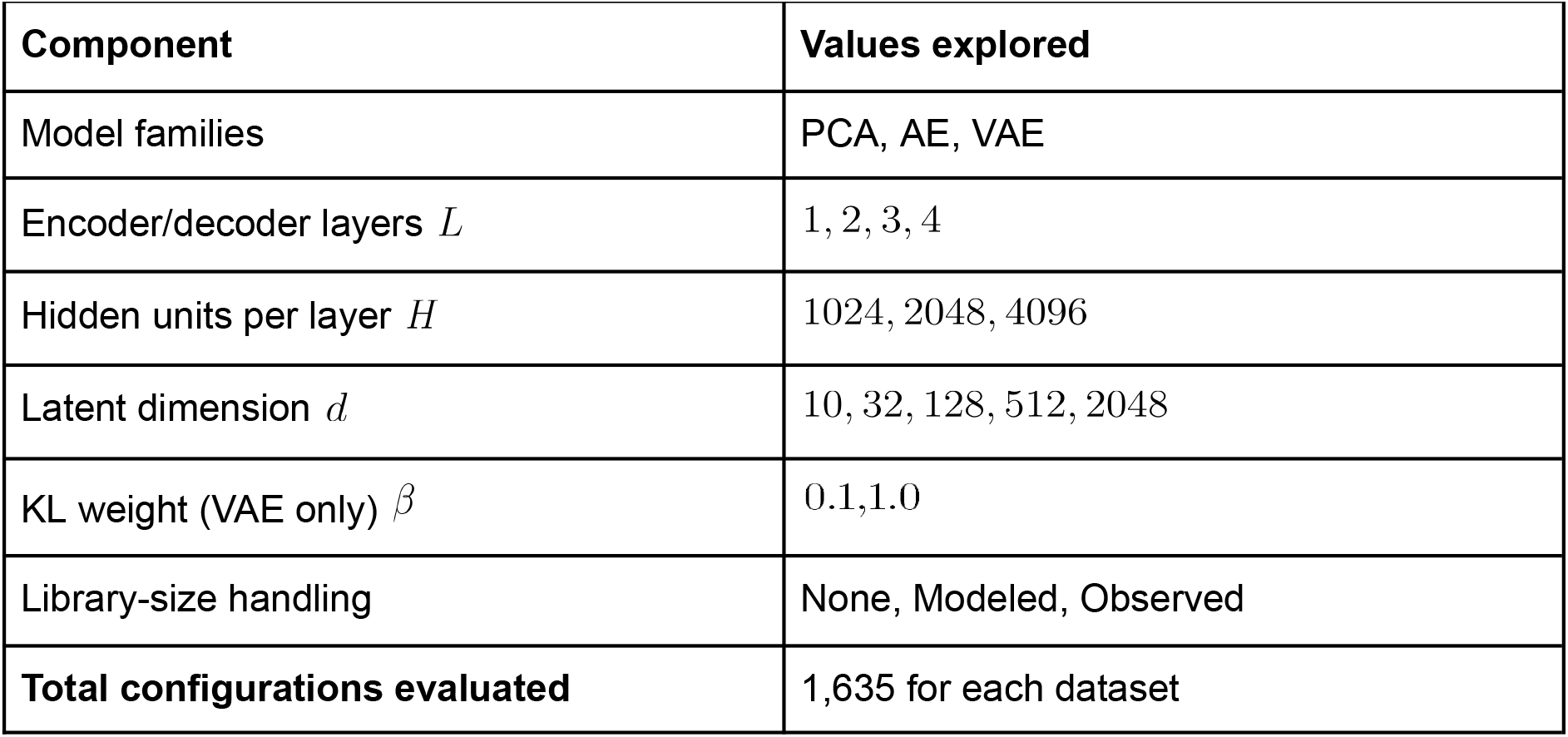
End-to-end reconstruction hyperparameter search space. We swept over MLP depth, width, latent dimension, library-size handling strategy, and (for VAE models) the KL weight. The best configuration was selected by reconstruction loss (MSE) on the validation set, and metrics were reported on the test set. PCA was evaluated only with library-size handling set to “None.”

**Supplementary Table 2.** Foundation-model reconstruction hyperparameter search space. The encoder is frozen (foundation model), and we sweep across decoder architecture (MLP, KNN, Transformer) and architecture-specific hyperparameters. For MLP decoders, we sweep depth and width matching the end-to-end search. For KNN decoders, we sweep the number of neighbors *k*. For Transformer decoders, we evaluate two pre-defined configurations (Small and Large). The best configuration was selected by reconstruction loss (MSE) on the validation set, and metrics were reported on the test set. Performance was largely insensitive to *k* within the tested range (Extended Data Fig. 7), and we used *k* = 10 for all main analyses.

| Component | Values explored |
| --- | --- |
| Frozen encoders | scGPT, SCimilarity, SE, scConcept |
| Decoder families | MLP, KNN, Transformer |
| MLP decoder layers $L$ | 1, 2, 3, 4 |
| MLP hidden units per layer $H$ | 1024, 2048, 4096 |
| KNN number of neighbors $k$ | 5, 10, 20, 50 |
| Transformer configurations | Small (4 heads, 3 layers); Large (8 heads, 6 layers) |

## Footnotes

1 STATE includes both a pretrained embedding model and a perturbation prediction model. For clarity, we use SE to refer to the embedding learned from the embedding model, and STATE to refer to the perturbation prediction model throughout.

